# Sodium channel toxin-resistance mutations do not govern batrachotoxin (BTX) autoresistance in poison birds and frogs

**DOI:** 10.1101/2020.10.29.361212

**Authors:** Fayal Abderemane-Ali, Nathan D. Rossen, Megan E. Kobiela, Robert A. Craig, Catherine E. Garrison, Lauren A. O’Connell, J. Du Bois, John P. Dumbacher, Daniel L. Minor

**Affiliations:** Cardiovascular Research Institute, University of California, San Francisco, California 93858-2330 USA; Departments of Biochemistry and Biophysics, and Cellular and Molecular Pharmacology, University of California, San Francisco, California 93858-2330 USA; California Institute for Quantitative Biomedical Research, University of California, San Francisco, California 93858-2330 USA; Kavli Institute for Fundamental Neuroscience, University of California, San Francisco, California 93858-2330 USA; Molecular Biophysics and Integrated Bio-imaging Division, Lawrence Berkeley National Laboratory, Berkeley, CA 94720 USA; School of Biological Sciences, University of Nebraska – Lincoln, Lincoln, Nebraska 68588; Department of Chemistry, Stanford University, Stanford, CA 94305; Department of Biology, Stanford University, Stanford, CA 94305; Institute for Biodiversity Science and Sustainability, California Academy of Sciences, San Francisco, CA 94118, USA; Department of Biology, San Francisco State University, San Francisco, CA 94132, USA

## Abstract

Poisonous organisms carry small molecule toxins that alter voltage-gated sodium channel (Na✓) function. Among these, batrachotoxin (BTX) from *Pitohui* toxic birds and *Phyllobates* poison frogs, stands out because of its lethality and unusual effects on Na_v_ function. How these toxin-bearing organisms avoid autointoxication remains poorly understood. In poison frogs, a Na_v_ DIVS6 pore-forming helix N→T mutation has been proposed as the BTX resistance mechanism. Here, we show that this variant is absent from *Pitohui* and poison frog Na_v_s, incurs a strong cost that compromises channel function, and fails to produce BTX-resistant channels when tested in the context of poison frog Na_v_s. We further show that captive-raised poison frogs are BTX resistant, even though they bear BTX-sensitive Na_v_s. Hence, our data refute the hypothesis that BTX autoresistance is rooted in Na_v_ mutations and instead suggest that more generalizable mechanisms such as toxin sequestration act to protect BTX-bearing species from autointoxication.

## Introduction

Many organisms harbor small molecule toxins that target ion channels as a means of defense from predation (1). Among these toxins, batrachotoxin (BTX) is a renowned steroidal amine that is found in distantly related vertebrate lineages including neotropical poison frogs *(Dendrobatidae)* (2) and multiple poisonous bird species (*Pitohui sp*. and *Ifrita kowaldi*) (3, 4). These animals acquire BTX from their diet (5–7) and accumulate it in their muscles as well as in their skin and feathers where this toxin is used as a predation defense (2, 8). BTX has an unusual mechanism of action against voltage-gated sodium channels (Na_v_s) that facilitates channel opening and prevents channel inactivation (9–12). How vertebrates that bear BTX or other small molecule toxins avoid autointoxication remains unresolved (13–15). Toxin-resistant mutants of target ion channels in the host organisms (16–19) or their predators (20–22) have been suggested as the primary driver of toxin autoresistance (2). *Phyllobates terribilis*, the poison frog carrying the highest BTX levels, is BTX-resistant, even when animals are captive-raised and lack toxin (23), suggesting that BTX resistance is a heritable trait. Partial sequencing of poison frog Na_v_s has suggested a number of Na_v_ mutations that might alter BTX sensitivity (24). One of these, a conserved Domain IV S6 helix (DIVS6) N→T variation in *P. terribilis* skeletal muscle Na_v_1.4, reduces BTX sensitivity when tested in rat Na_v_1.4 (25). This result has been used to support the notion that BTX-resistance relies on toxin-resistant channel mutants, similar to examples for tetrodotoxin (TTX) (18–22) and saxitoxin (STX) (17) resistant Na_v_s, and epibatidine-resistant nicotinic acetylcholine receptors (nAChRs) (16). However, this DIVS6 N→T change occurs with very low frequency among wild *P. terribilis* (26) and is absent in wild *Phyllobates aurotaenia* (24, 26), another poison frog species known to carry high BTX levels (27). Further, in the wild there are a number of examples of poisonous frogs (24, 26) and predators of toxic animals (28) that lack toxin-resistant mutations, raising questions about generality of the target-based mechanism. Given these issues and the fact that no functional studies of poison frog Na_v_s have been published, whether BTX-bearing poison animals rely on Na_v_ mutations or some other mechanism for BTX autoresistance remains unresolved.

Here, we cloned and characterized Na_v_s from the BTX-bearing bird *Pitohui uropygialis meridionalis (Pum)* and two poison frog species that carry alkaloid toxins in the wild *(Phyllobates terribilis*, BTX, and *Dendrobates tinctorius*, histrionicotoxin HTX, and pumiliotoxin, PTX). Surprisingly, we did not find the DIVS6 N→T mutation in any of these species even though the poison frog Na_v_s possessed all other mutations suggested to contribute to BTX resistance (24). Further, the *Pitohui* and poison frog Na_v_s were BTX sensitive. Although the DIVS6 N→T mutation reduced *Pum* Na_v_ BTX sensitivity, in line with its effect in rat Na_v_1.4 (25), this same mutation failed to confer BTX resistance to poison frog Na_v_s. Additionally, in all cases, DIVS6 N→T greatly compromised channel function, underscoring the tradeoffs between toxin-resistant mutations and fitness cost (29). Most surprising, poison frogs having BTX-sensitive Na_v_s proved resistant to BTX poisoning and resisted poisoning by another small molecule toxin, saxitoxin (STX), despite expressing STX-sensitive Na_v_s. A distantly related poison frog that does not carry BTX *(Mantella)*also proved resistant to BTX and STX. Hence, our studies refute the idea that target Na_v_ mutations are singularly responsible for BTX autoresistance in poisonous birds and frogs, and suggest that other, more general mechanisms such as toxin sequestration (13, 14, 30, 31) defend against autointoxication.

## Results

### Cloning and characterization of skeletal and cardiac *Pitohui* Na_v_s

*Pitohui* is one of only a few bird genera known to carry BTX (3, 4, 32) and has BTX levels in its skeletal and cardiac muscles that should alter Na_v_ function (8). To investigate possible mechanisms of BTX resistance, we used a *Pitohui* genomic DNA library to identify and assemble genes for two *Pitohui uropygialis meridionalis* Na_v_s, skeletal muscle *Pum* Na_v_1.4 (Fig. S1) and cardiac *Pum* Na_v_1.5 (Fig. S2). Primary sequence alignment showed extensive similarities between *Pum* Na_v_1.4, *Pum* Na_v_1.5, and other vertebrate homologs (~73% amino acid identity) (Figs. S1 and S2), including hallmark Na_v_ features, such as: a selectivity filter ‘DEKA’ motif, canonical RXXR repeats in S4 in all four voltage sensor domains, and the ‘IFM’ motif responsible for fast inactivation (33) (Figs. S1 and S2).

Whole-cell patch clamp electrophysiology of *Pum* Na_v_1.4 and *Pum* Na_v_1.5 transfected into HEK293 cells demonstrated that both have fast voltage-dependent activation followed by a fast and complete voltage-dependent inactivation typical of Na_v_s (Figs. 1a-b, S3a-b, and Tables 1 and S1) similar to human Na_v_1.4, *Hs* Na_v_1.4, recorded under identical conditions (Fig. 1c-d, Tables 1 and S1). Because Na_v_s can harbor resistance mutations to other small molecule toxins (13, 14, 17, 20), we anticipated that the *Pitohui* Na_v_s might be BTX-resistant. Surprisingly, application of 10 μM BTX drastically altered the function of both *Pum* Na_v_s, yielding typical BTX-induced functional consequences: a hyperpolarized shift in the voltage-dependency of activation (ΔV_1/2 BTX_ = −33.6 ± 1.2 and −37.4 ± 1.8 for *Pum* Na_v_1.4 and *Pum* Na_v_1.5, respectively), reduced inactivation, and enhanced tail currents (10, 11) (Figs. 1a-b and S3a-b, Table 1). The BTX-induced activation curve follows a double Boltzmann function in which the first and second components arise from BTX-bound and unmodified channels, respectively (34). Notably, the BTX-induced changes were equivalent to those elicited by BTX application to *Hs* Na_v_1.4 (ΔV_1/2 BTX_ = −35.9 ± 1.8) (Fig. 1d, Table 1). These data demonstrate that despite the fact that *Pitohui* carry BTX in its skeletal muscles and heart (8), their skeletal and cardiac Na_v_s are BTX-sensitive. Thus, autoresistance cannot originate from altered BTX sensitivity in the two most likely target channels exposed to lethal BTX levels.

**Figure 1.**
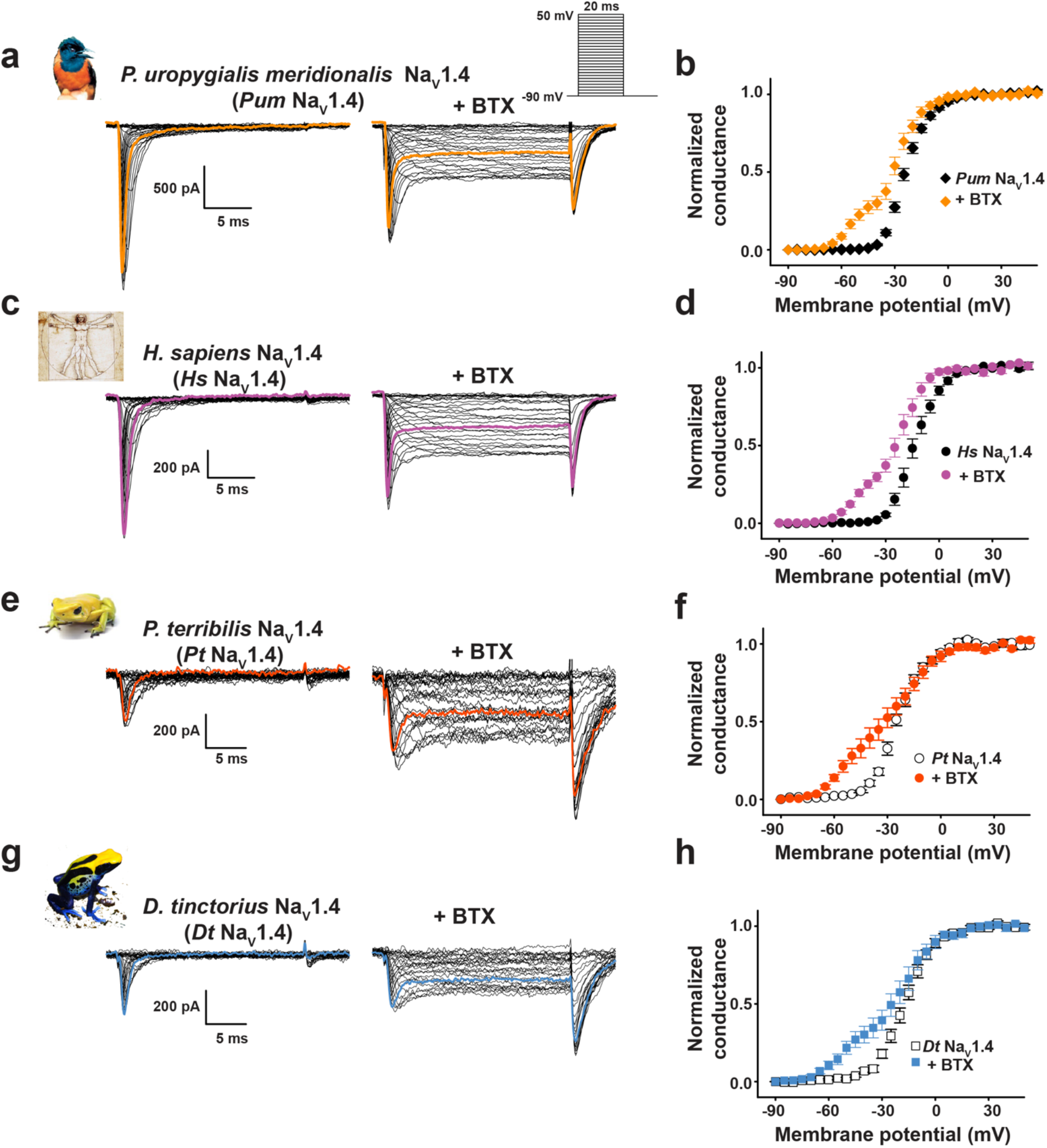
*Pitohui* and Poison frog Na_v_1.4 channels are BTX sensitive. Exemplar current recordings (**a**, **c**, **e**, and **g**) for: **a**, *Pitohui uropygialis meridionalis* Na_v_1.4 *(Pum* Na_v_1.4); **c**, Human Na_v_1.4 (*Hs* Na_v_1.4); **e**, *Phyllobates terribilis* Na_v_1.4 (*Pt* Na_v_1.4); and **g**, *Dendrobates tinctorius* Na_v_1.4 (*Dt* Na_v_1.4) expressed in HEK293 cells in absence (left) or presence of 10 μM BTX (right). Trace at 0 mV is highlighted in each panel. Currents were evoked with the shown multistep depolarization protocol (inset panel ‘**a**’). Conductance-voltage relationships (**b**, **d**, **f**, and **h**) in presence or absence of 10 μM BTX for **b**, *Pum* Na_v_1.4 (black diamonds), +BTX (orange diamonds), **d**, *Hs* Na_v_1.4 (black circles), +BTX (purple circles) **f**, *Pt* Na_v_1.4 (white circles), +BTX (orange circles), and **h**, *Dt* Na_v_1.4 (white squares), +BTX (blue squares).

**Table 1.**
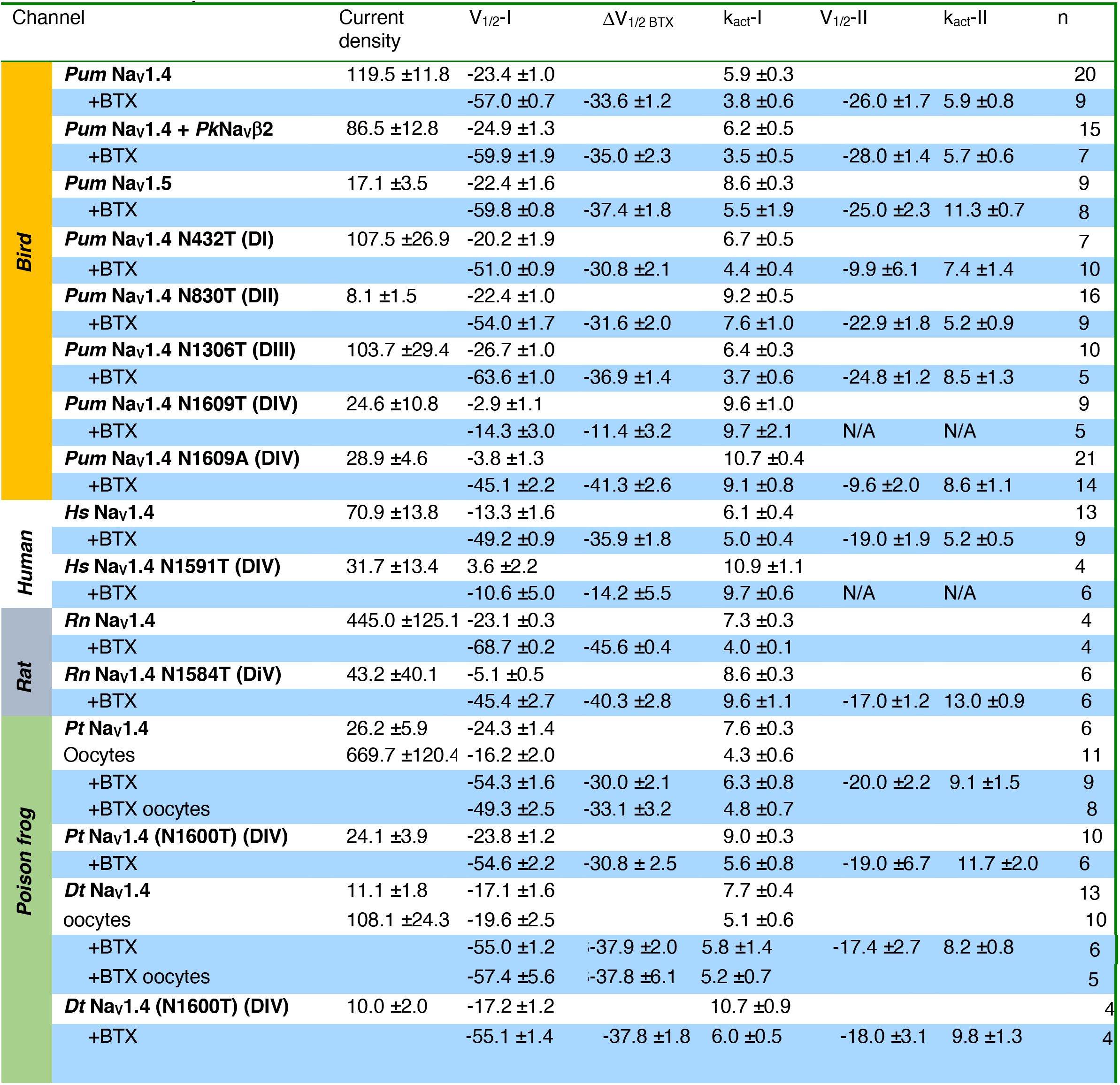
Activation parameters

Na_v_s are often co-expressed with auxiliary β subunits that can alter their biophysical (35) and pharmacological (36, 37) properties. To test whether this subunit could impact BTX resistance, we identified the *Pitohui uropygialis meridionalis* gene encoding for a transmembrane protein bearing the key features of Na_v_β2, including a pair of conserved disulfides (38). (Fig. S3c). Co-expression of *Pum* Na_v_β2 with *Pum* Na_v_1.4 had no impact on channel biophysical properties or on BTX responses relative to *Pum* Na_v_1.4 alone (Fig. S3d-e, Tables 1 and S1). Thus, *Pum* Na_v_1.4 alone and *Pum* Na_v_1.4 in combination with *Pum* Na_v_β2 failed to showed evidence of BTX-resistant channels.

### Cloning and characterization of poison frog Na_v_s

Poison frogs in the genus *Phyllobates* (Family *Dendrobatidae)* are the most well-known BTX carriers (2, 27, 39). A number of studies have identified amino acid substitutions hypothesized to contribute to poison frog Na_v_ BTX resistance (24–26). However, cloning and functional characterization of poison frog Na_v_s is noticeably absent from the literature. We used skeletal muscle from captive-raised members of two representative poison frog species, one that carries high levels of BTX in the wild, *Phyllobates terribilis (Pt* Na_v_1.4) (23, 24, 40) and one not known to carry BTX, *Dendrobates tinctorius (Dt* Na_v_1.4)(24), to clone poison frog Na_v_1.4s. Consistent with evolutionary relationships between the two species (24, 26), *Pt* Na_v_1.4 and *Dt* Na_v_1.4 were highly similar to each other (~95% amino acid identity) and other vertebrate Na_v_s (~73% amino acid identity), and bore all Na_v_ hallmarks (Fig. S1). Importantly, their DIS6 and DIVS6 sequences were identical to those reported previously (24) with the remarkable absence in *Pt* Na_v_1.4 of the proposed BTX-resistance mutation DIVS6 N→T (*Pt* Na_v_1.4 Asn1600, *Pt* Na_v_1.4 N1584T (rat numbering) (25)) (Fig. S1, Table S2). Genomic DNA sequencing covering the *Pt* Na_v_1.4 DIVS6 yielded nucleotide sequences identical to those obtained from cDNA and cross-validated the absence of the DIVS6 N→T substitution. These findings are consistent with the observation that the DIVS6 N1600T substitution has a very low frequency among wild *P. terribilis* (26). Besides the prior reported amino acid variants (24), *Pt* Na_v_1.4 and *Dt* Na_v_1.4 had 93 additional positions distributed throughout the channel that differed from conserved bird, human, and rat Na_v_1.4 residues (Fig. S4, Table S2). Although we could readily sequence the *Pt* Na_v_1.4 and *Dt* Na_v_1.4 genes, both proved prone to recombination and deletion upon passage through *E. coli*, rendering the native DNA sequences impossible to handle. To solve this problem, we re-designed the codon usage to preserve the amino acid sequence of both. These re-designed genes were well behaved and allowed us to conduct electrophysiological characterization of *Pt* Na_v_1.4 and *Dt* Na_v_1.4 in mammalian and amphibian expression systems.

Whole-cell patch clamp electrophysiology of HEK293 cells transfected with *Pt* Na_v_1.4 and *Dt* Na_v_1.4 yielded voltage-dependent channels that matched the properties of both *Pum* and *Hs* Na_v_1.4s (Fig. 1e-h, Tables 1 and S1). Strikingly, both poison frog Na_v_s had the same response as *Pum* and *Hs* Na_v_1.4 to 10 μM BTX (Fig. 1e-h), (ΔV_1/2 BTX_= −30.0 ± 2.1 and −37.9 ± 2.0 mV for *Pt* Na_v_1.4 and *Dt* Na_v_1.4, respectively), demonstrating that these poison frog channels are not resistant to BTX and ruling out the possibility that the >90 amino acid variants between poison frog and human channels, including the previously proposed changes in DIS6 and DIVS6 (24), could cause BTX-resistance.

Because expression of amphibian channels in an amphibian cell could provide a more native-like context, we also expressed *Pt* Na_v_1.4 and *Dt* Na_v_1.4 in *Xenopus* oocytes and examined their function by two-electrode voltage clamp. Both channels had biophysical parameters that matched those measured in mammalian cells (Fig. S5a-d, Table 1). Further, BTX application caused the strong hallmark functional modification observed for all of the other channels we studied (Figs 1, S3a-b and d-e, and S5a-d), including voltage-dependent activation shifts comparable to those measured in mammalian cells (ΔV_1/2 BTX_ = −33.1 ± 3.2 and −30.0 ± 2.1; and −37.8 ± 6.1 and −37.9 ± 2.0 mV for *Pt* Na_v_1.4 and *Dt* Na_v_1.4 expressed in *Xenopus* oocytes and HEK293 cells, respectively). The shift was more complete in oocytes (*cf*. Figs. 1f and h and S5b and d), a result that likely originates from the fact that for technical reasons BTX was injected into the oocytes rather than applied by bath application as for mammalian cells. Taken together, the biophysical characterization of *Pt* Na_v_1.4 and *Dt* Na_v_1.4 demonstrates that these channels are not BTX-resistant. Our combined observations that Na_v_s from two classes of BTX-carrying animals, *Pitohui* and *P. terribilis*, are vulnerable to BTX challenge the idea that Na_v_ mutation is the BTX autoresistance strategy as suggested for poison frogs such as *P. terribilis* (24, 25).

### DIVS6 N→T mutation fails to confer BTX resistance to poison frog Na_v_s

Because the DIVS6 N→T mutation was absent from Na_v_s of BTX-bearing species, we wondered whether the observation that DIVS6 N→T could confer BTX resistance to rat Na_v_1.4 (25) was impacted by the >90 amino acid differences between poison frog and mammalian Na_v_s (Fig. S4, Table S2). Therefore, we placed the DIVS6 N→T mutation in poison bird, human, and poison frog Na_v_1.4s (*Pum* Na_v_1.4 N1609T, *Hs* Na_v_1.4 N1591T, *Pt* Na_v_1.4 N1600T, and *Dt* Na_v_1.4 N6100T) and measured its effects on channel function and BTX sensitivity. Consistent with studies of rat Na_v_1.4 DIVS6 N→T (25), DIVS6 N→T eliminated the ability of BTX to block inactivation and induce large tail currents in *Pum* Na_v_1.4 and *Hs* Na_v_1.4 (Fig. 2a-d). Nevertheless, the bird and human Na_v_1.4s were not rendered completely BTX resistant. Application of 10 μM BTX shifted the voltage-dependent activation of both channels, making them more easily opened by voltage (ΔV_1/2 BTX_ = −11.4 ± 3.2 and −14.2 ± 5.5 mV for *Pum* Na_v_1.4 N1609T and *Hs* Na_v_1.4 N1591T, respectively) (Fig. 2b and d, Table 1). Further, the BTX-induced double Boltzmann was lost (Fig. 2b and d), suggesting an enhanced BTX affinity. Due to its limited effectiveness in blocking the effects of BTX in the bird and human channels, we revisited the consequences of the DIVS6 N→T mutation in rat Na_v_1.4 *(Rn* Na_v_1.4). Similar to the results with bird and human channels, DIVS6 N→T reduced but did not eliminate *Rn* Na_v_1.4 channel BTX sensitivity (Figure S6a-d). Application of 10 μM BTX shifted the voltage-dependent activation of both *Rn* Na_v_1.4 and *Rn* Na_v_1.4 N1584T making them more easily opened by voltage (ΔV_1/2 BTX_ = −45.6 ± 0.4 and −40.3 ± 2.8 mV for *Rn* Na_v_1.4 and *Rn* Na_v_1.4 N1584T, respectively) (Fig S6b and d, Table 1). Thus, DIVS6 N→T was unable to mitigate the effects of BTX completely in any Na_v_ orthologues. In all three contexts, DIVS6 N→T also affected channel biophysical properties (Figs. S6e-g, S7a-h, Tables 1, S1 and S3). DIVS6 N→T rendered *Pum* Na_v_1.4, *Hs* Na_v_1.4 and *Rn* Na_v_1.4 more difficult to open, shifting the activation voltage dependence to depolarizing potentials (ΔV_1/2_ =+20.5 ± 1.5, +16.9 ± 2.7, and +18.0 ± 0.6 mV for *Pum* Na_v_1.4 N1609T, *Hs* Na_v_1.4 N1591T, and *Rn* Na_v_1.4 N1584T, respectively) (Figs. S6e, S7b and f, Tables 1 and S3), and made the channels easier to inactivate, shifting the voltage dependence of steady state inactivation towards hyperpolarizing potentials (ΔV_1/2 inact_ = −10.3 ± 2.1, −9.8 ± 1.2 and −27.0 ± 0.4 mV for *Pum* Na_v_1.4 N1609T, *Hs* Na_v_1.4 N1591T and *Rn* Na_v_1.4 N1584T, respectively) (Figs. S6f, S7c and g, Tables S1 and S3). Further, DIVS6 N→T diminished *Pum* Na_v_1.4 N1609T, *Hs* Na_v_1.4 N1591T and *Rn* Na_v_1.4 N1584T current densities by 79%, 55% and 90%, respectively (Figs. S6a, c, and g, and S7a, d, e, and h, Tables 1 and S3). Thus, the DIVS6 N→T change incurs a substantial functional cost.

**Figure 2.**
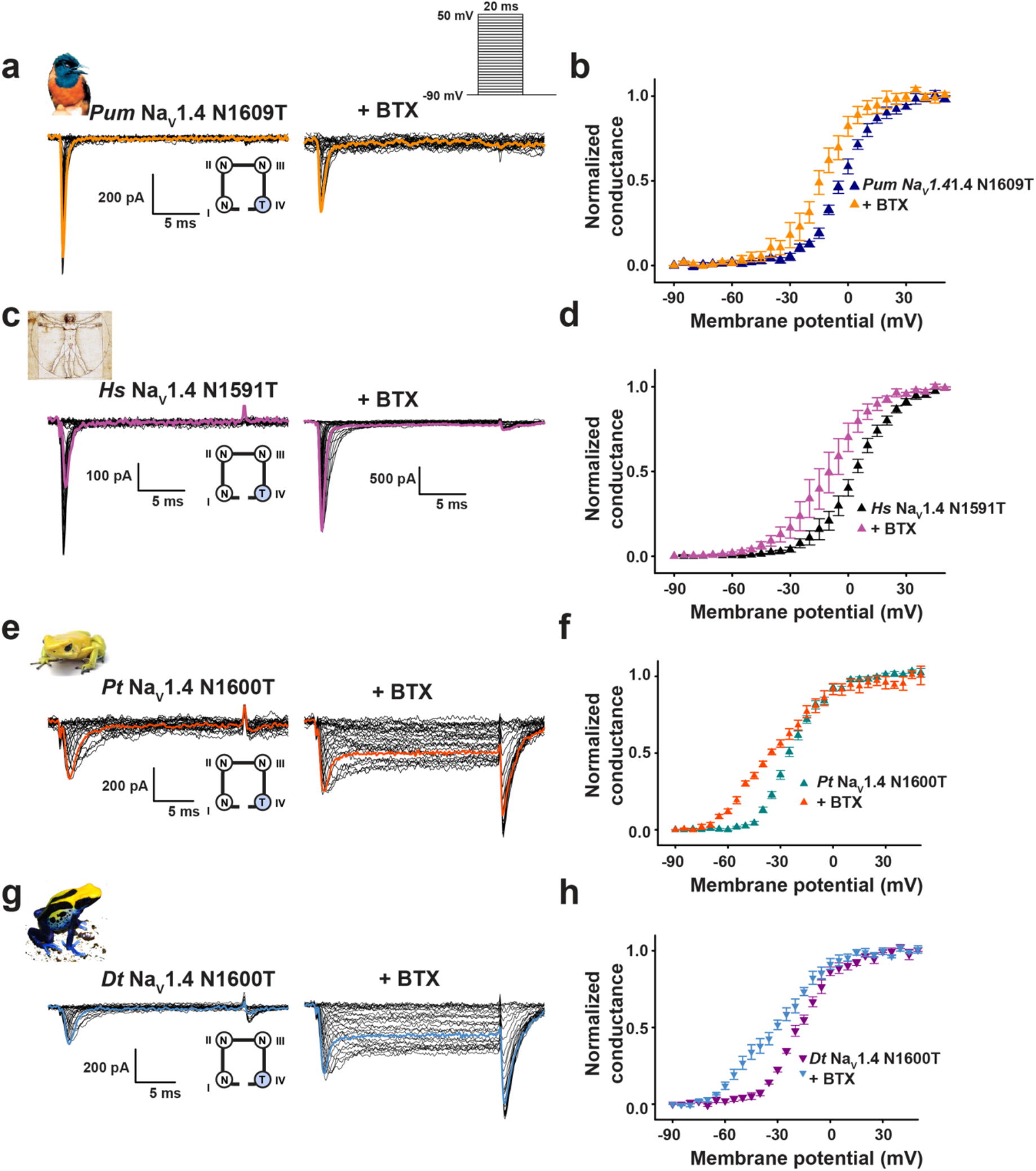
DIVS6 N→T mutation reduces BTX sensitivity of *Pitohui* and human but not poison frog Na_v_1.4s. Exemplar current recordings (**a**, **c**, **e**, and **g**) for: **a**, *Pum* Na_v_1.4 N1609T; **c**, *Hs* Na_v_1.4 N1591T; **e**, *Pt* Na_v_1.4 N1600T; and **g**, *Dt* Na_v_1.4 N1600T expressed in HEK293 cells in absence (left) or presence of 10 μM BTX (right). Trace at 0 mV is highlighted in each panel. Currents were evoked with the shown multistep depolarization protocol (inset panel ‘**a**’). Cartoon shows a diagram of the identities of the S6 ‘Asn’ for each construct. Conductance-voltage relationships (**b**, **d**, **f**, and **h**) for **b**, *Pum* Na_v_1.4 N1609T, **d**, *Hs* Na_v_1.4 N1591T **f**, *Pt* Na_v_1.4 N1600T, and **h**, *Dt* Na_v_1.4 N1600T in presence or absence of 10 μM BTX.

To probe the DIVS6 Asn site further, we examined the consequences of mutation to alanine in *Pum* Na_v_1.4. *Pum* Na_v_1.4 N1609A phenocopied the biophysical changes measured for N1609T producing channels that were more difficult to open (ΔV_1/2_ =+19.6 ± 1.6), easier to inactivate (ΔV_1/2 inact_ = −13.4 ± 1.6), and had current density reduced by 76% (Fig. S7a-d, Tables 1, S1, and S3), in agreement with the reduced channel activity reported for the corresponding rat Na_v_1.4 mutant (41, 42). These biophysical changes match those of the BTX-resistant *Pum* Na_v_1.4 N1609T; however, *Pum* Na_v_1.4 N1609A retained all of the classical BTX responses such as reduction of inactivation, enhanced tail current, and a leftward shift of the activation voltage dependence (Fig. S7i-j, Tables 1 and S3). The failure of the N1609A to diminish BTX sensitivity shows that the reduction of BTX sensitivity in *Pum* Na_v_1.4 N1609T, *Hs* Na_v_1.4 N1591T, and *Rn* Na_v_1.4 N1584T is a specific effect of the threonine mutation, and not a consequence of the changes in channel biophysical properties or current density reduction (Table S3).

To our surprise, placing DIVS6 N→T in both poison frog Na_v_1.4s failed to blunt the effects of BTX on channel activation and inactivation (Fig. 2e-h) (ΔV_1/2 BTX_ = −30.0 ± 2.1, −30.8 ± 2.5 mV for *Pt* Na_v_1.4 and *Pt* Na_v_1.4 N1600T, respectively and −37.9 ± 2.0 and −37.8 ± 1.8 for *Dt* Na_v_1.4 and *Dt* Na_v_1.4 N1600T, respectively). Hence, even though the DIVS6 N→T change reduces *Pum* Na_v_1.4, *Hs* Na_v_1.4, and *Rn* Na_v_1.4 BTX responses (Figs. 2a-d, S6a-d), this same change fails to affect the BTX sensitivity of poison frog Na_v_s. Unlike its effects in *Pum, Hs*, and *Rn* Na_v_s, DIVS6 N→T did not cause major changes in poison frog Na_v_ biophysical properties, shifting only the inactivation voltage dependence by −10 mV while leaving the activation voltage dependence and current density unchanged (Fig. S8a-c, Tables 1, S1, and S3). However, expression of *Pt* Na_v_1.4 N1600T and *Dt* Na_v_1.4 N1600T in *Xenopus* oocytes revealed dramatic reductions in channel activity (Fig. S8d-h) that prevented measurement of channel biophysical properties and BTX responses. These results reveal that DIVS6 N→T is detrimental to function and may interfere with channel folding and maturation in a manner that is accentuated at lower temperatures, such as those used to store incubate the oocytes. This context-dependent loss of function indicates that the DIVS6 N→T variant exacts a functional cost that, together with its ineffectiveness in endowing poison frog Na_v_s with BTX resistance, refutes the idea that DIVS6 N→T could serve as an effective BTX autoresistance mechanism.

### Cost of the conserved N→T mutation is context dependent

The varied outcomes of DIVS6 N→T on BTX sensitivity among the poison bird, human, rat, and poison frog Na_v_s highlight the importance of context in determining the functional consequences of mutations. Because the equivalent residue is conserved in all four S6 helices (Figs. S1 and S2), we systematically introduced S6 N→T into each of the *Pum* Na_v_1.4 S6 segments and measured channel properties and BTX responses to investigate the question of context dependent effects further (Fig. 3). Whole cell patch clamp recordings from HEK293 cells transfected with these mutant channels revealed clear, domain-specific differences. Contrasting the effect of DIVS6 N→T (Fig. S7b), voltage-dependent activation of channels having the N→T mutation in DI, DII, or DIII (*Pum* Na_v_1.4 N432T (DI), *Pum* Na_v_1.4 N830T (DII), and *Pum* Na_v_1.4 N1306T (DIII)) was unchanged relative to wild-type (V_1/2_ = −20.2 ± 1.9, −22.4 ± 1.0, −26.7 ± 1.0, and −23.4 ± 1.0 mV for *Pum* Na_v_1.4 N432T (DI), *Pum* Na_v_1.4 N830T (DII), *Pum* Na_v_1.4 N1306T (DIII), and *Pum* Na_v_1.4, respectively) (Figs. 3a-j, S9a-b, and Tables 1 and S3). By contrast, we found varied effects on steady state inactivation. DI and DIII changes showed wild-type like behavior, whereas the DII mutant had a ~10 mV hyperpolarizing shift (V_1/2 inact_ = −61.2 ± 1.6, −75.0 ± 0.9, −65.3 ± 1.0, and −64.2 ± 1.3 mV for *Pum* Na_v_1.4 N432T (DI), *Pum* Na_v_1.4 N830T (DII), *Pum* Na_v_1.4 N1306T (DIII), and *Pum* Na_v_1.4, respectively) (Fig. S9c, Tables S1 and S3). All three had strong BTX responses similar to wild-type (ΔV_1/2 BTX_ = −30.8 ± 2.1, −31.6 ± 2.0, −36.9 ± 1.4, and −33.6 ± 1.2 mV for *Pum* Na_v_1.4 N432T (DI), *Pum* Na_v_1.4 N830T (DII), *Pum* Na_v_1.4 N1306T (DIII), and *Pum* Na_v_1.4, respectively) (Fig. 3, Tables 1 and S3). Thus, the only site where the conserved S6 N→T change affects BTX responses is in DIVS6, in line with its proposed contribution to the BTX binding site (25).

**Figure 3.**
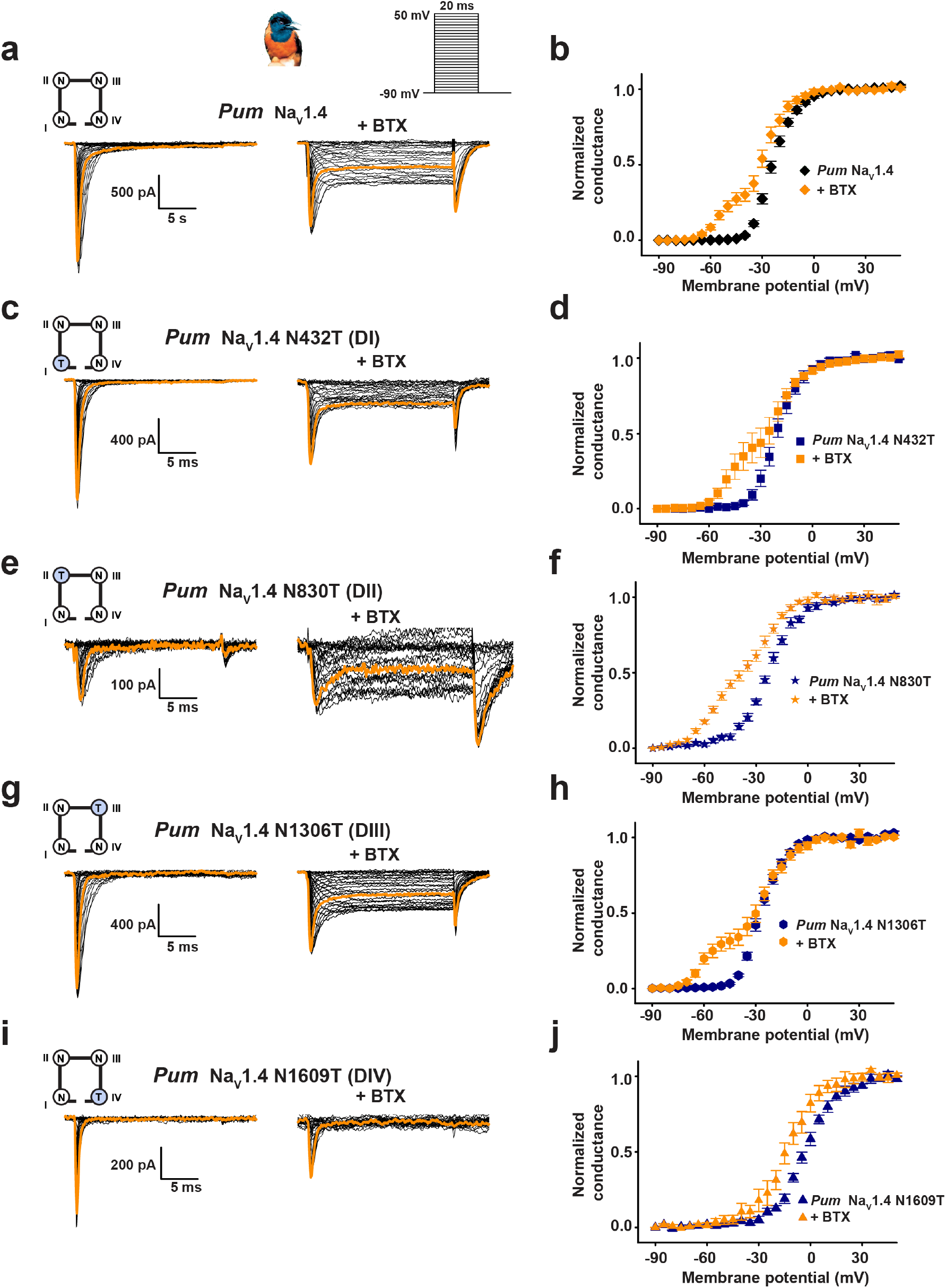
Sodium channel modulation by BTX is associated with an asymmetry at the inner pore level. Exemplar current recordings (**a**, **c**, **e**, **g**, and **i**) for: **a**, *Pum* Na_v_1.4; **c**, *Pum* Na_v_1.4 N432T; **e**, *Pum* Na_v_1.4 N830T; **g**, *Pum* Na_v_1.4 N1306T, and **i**, *Pum* Na_v_1.4 N1609T expressed in HEK293 cells in absence (left) of presence of 10 μM BTX (right). Trace at 0 mV is highlighted in each panel. Currents were evoked with the shown multistep depolarization protocol (inset panel ‘**a**’). Cartoon shows a diagram of the identities of the S6 ‘Asn’ for each construct. Conductance-voltage relationships (**b**, **d**, **f**, **h**, and **j**) in presence or absence of 10 μM BTX for **b**, *Pum* Na_v_1.4 (black diamonds), +BTX (orange diamonds) **d**, *Pum* Na_v_1.4 N432T (dark blue squares), +BTX (orange squares), **f**, *Pum* Na_v_1.4 N830T (dark blue stars), +BTX (orange stars), and **h**, *Pum* Na_v_1.4 N1306T (dark blue hexagons), +BTX (orange hexagons), and **j**, *Pum* Na_v_1.4 N1609T (dark blue triangles), +BTX (orange triangles). Data in ‘**a**’ and ‘**b**’ are from Figure 1 **a** and **b**.

As with the biophysical changes, the effects on current density from placing the N→T change in different channel domains were not uniform. The DIS6 and DIIIS6 N→T mutants had current densities matching wild-type (Figs. 3a, c, g, S9a and d, and Tables 1 and S3), whereas, DIIS6 N→T lowered the current density and was more detrimental to channel activity than DIVS6 N1609T or N1609A (Figs. 3a, e, S9a and d, and Tables 1 and S3). Together, these data show that there is no correlation between changes in channel biophysical properties and the acquisition of BTX resistance and are in line with the results from DIVS6 N→T and N→A mutants (Figs. 2, S7, and S8, Table S3).

Consideration of the conserved S6 asparagine structural context provides insight into the context-dependent effects. The two S6 sites where N→T has no impact on channel biophysics, BTX responses, or current density, DIS6 and DIIIS6, occupy positions that are partially exposed to the channel inner pore (Fig. S9e-f). By contrast, the two positions that affect channel biophysics and current density, DIIS6 and DIVS6, interact with the S4-S5 linkers (43) (Fig. S9e-f) and altering these buried sites comes with substantial functional costs. Hence, DIVS6 N→T carries major disadvantages for protecting animals such as *Pitohui* and poison frogs against BTX autointoxication.

### Toxin-free poison frogs are BTX resistant in the absence of the DIVS6 N→T mutation

The surprising observation that Na_v_s from BTX-carrying birds and frogs remain BTX-sensitive raised the question of whether the species from which we cloned the channels were actually BTX resistant. Because of difficulties in obtaining live animals, we were unable to investigate *Pitohui*BTX resistance. Captive-raised poison frogs lack BTX, as this toxin is acquired in the wild from their diet (5, 6). Thus, it was possible that the toxin-free poison frogs used to clone Na_v_s were not BTX resistant due to the absence of selective pressure from the toxin, a possibility underscored by the high functional cost of DIVS6 N→T (Figs. 2e-h and S8, Tables 1, S1 and S3). To test whether captive-raised poison frogs were BTX resistant, we conducted a series of toxin challenge experiments using five different frog species: two non-poisonous frogs *(Xenopus laevis* and *Polypedates leucomystax)*, two captive-raised poison frogs that carry alkaloid toxins in the wild (*Phyllobates terribilis*, BTX; *Dendrobates tinctorius*, histrionicotoxins HTX, and pumiliotoxins, PTX), and an unrelated captive-raised Malagasy poison frog that carries PTX rather than BTX in the wild (*Mantella aurantiaca*) and represents an independent evolutionary origin of chemical defenses (44–46). We challenged these animals with three different toxins that target Na_v_s: BTX, and two guanidinium toxins that act by a pore-blocking mechanism, saxitoxin (STX) (47, 48), and tetrodotoxin (TTX) (48).

We assessed the duration of recovery from anesthesia-induced paralysis following intramuscular injection of each toxin at 20 times the lethal dose based on values for mice (LD_50_) by monitoring how long it took the frog to show clear motor activity relative to injection of a phospho-buffered saline (PBS) control. Following BTX injection, *Xenopus laevis* and *Polypedates leucomystax* displayed an accelerated recovery from anesthesia that was at least two times faster than PBS injections (PBS and BTX recovery times: 29 ± 1 min. and 15 ± 5 min., and 169 ± 12 min. and 70 ± 20 min. for *X. laevis* and *P. leucomystax*, respectively) (Fig. 4a-b and f, Table S4). After the initial recovery, BTX was ultimately lethal to *X. laevis* (Fig.4a and f). By contrast, BTX injection did not change the anesthesia recovery time or kill any of the poison frogs, regardless of whether they carry BTX in the wild environment (*P. terribilis)* or are naturally BTX-free but harbor other alkaloid toxins (*D. tinctorius, M. aurantiaca)* (Fig. 4c-f, Table S4). Reponses to STX also revealed differences between non-poisonous and poison frogs. STX injection was lethal to *X. laevis* and *P. leucomystax* (Fig. 4a-b and f), whereas all three poison frogs fully recovered from anesthesia after STX injections (Fig. 4c-f). TTX was lethal to *X. laevis* (Fig. 4a and f). Although TTX caused extended paralysis in all other tested frogs, it was not lethal (Fig. 4a-f, Table S4). Thus, all tested poisonous frogs showed resistance to all three toxins while non-poisonous frogs were vulnerable to either BTX and STX (*P. leucomystax)* or all three toxins (*X. laevis)* (Fig. 4f).

**Figure 4.**
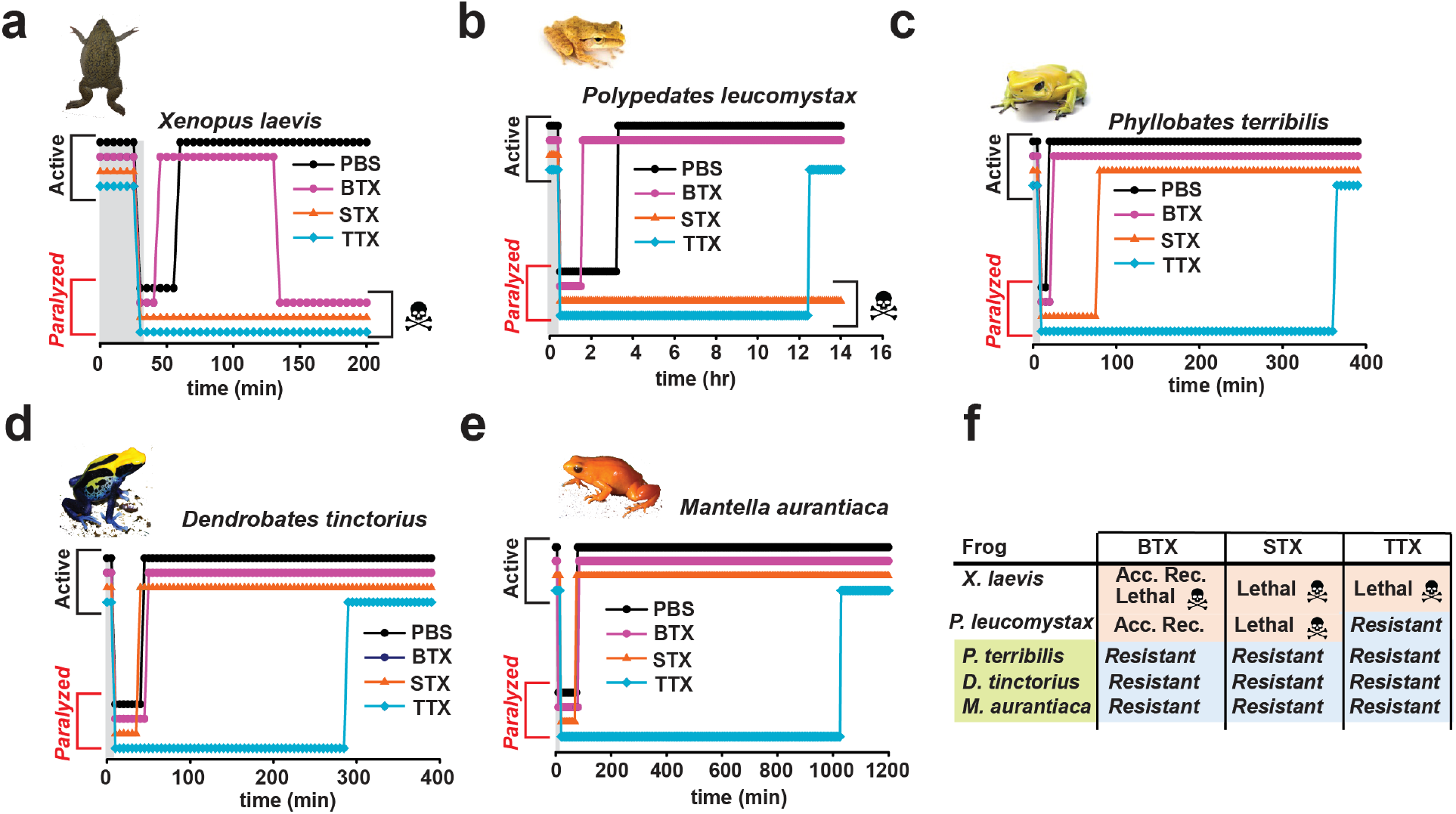
Captive-raised poison frogs are resistant to BTX and STX. **a-f**, Challenge experiments for **a**, *Xenopus laevis*,**b**, *Polypedates leucomystax*,**c**, *Phyllobates terribilis*,**d**, *Dendrobates tinctorius*, and **e**, *Mantella aurantiaca* to PBS (black circles), BTX (magenta circles), STX (orange triangles), or TTX (cyan diamond) injection. Grey area shows period of anesthesia application. Active and paralyzed states of the frogs are indicated. **f**,Summary of the sensitivity of the indicated species to BTX, STX and TTX. ‘Acc. Rec.’ denotes accelerated recovery from anesthesia. ‘Resistant’ denotes no toxin-induced death.

The striking differences in BTX-induced accelerated recovery from anesthesia between the non-poisonous and poisonous species was unexpected. Studies of eukaryotic and prokaryotic Na_v_s suggest that BTX and local anesthetics, such as the tricaine used for frog anesthesia, have overlapping binding sites within the channel pore (49–52). We considered that the differences in BTX-dependent accelerated recovery from anesthesia were a physiological manifestation of this molecular competition and indicated that BTX was engaging the target channels in non-poisonous frogs but not in poison frogs. Hence, we tested whether tricaine and BTX produced competing effects on *Pt* Na_v_1.4 and *Dt* Na_v_1. 4. Consistent with its anesthetic effects on the frogs, 0.5 mM tricaine inhibited both poison frog Na_v_s and had similar effects on the *Pum* Na_v_1.4 control (Fig. S10a-c and g). Subsequent BTX injection into the same tricaine-treated oocytes caused a complete relief of the tricaine block (Fig. S10a-c and g), in line with a direct competition between tricaine and BTX. By contrast, in the absence of tricaine, this dramatic BTX-induced increase in peak current was absent (Fig. S10d-f and h). These data demonstrate that the poison frog Na_v_s are competent for BTX-tricaine competition. Hence, the differences in BTX-induced accelerated recovery from anesthesia reflect the direct competition of the two compounds on the channel in the non-poisonous frogs and suggest that the poison frogs have a means to prevent BTX form engaging their Na_v_s.

The poison frog STX resistance we observed could be explained by a lack of Na_v_ sensitivity to this toxin. To test this possibility, we compared the responses of *Pt* Na_v_1.4, *Dt* Na_v_1.4, and *Pum* Na_v_1.4 as a control, to STX and TTX as the former had minimal effect on the poison frogs, whereas the latter caused potent paralysis (Fig. 4, Table S4). Extracellular application of increasing STX concentrations inhibited all three Na_v_s with a nanomolar response that matched other Na_v_s (47, 53, 54) (IC_50_ = 12.6 ± 1.4 nM, 14.6 ± 0.6 nM and 7.3 ± 0.5 nM for *Pt* Na_v_1.4, *Dt* Na_v_1.4 and *Pum* Na_v_1.4, respectively) (Fig. S11a-d). *Pt* Na_v_1.4 and *Dt* Na_v_1.4, and the control *Pum* Na_v_1.4 also had nanomolar TTX responses, similar to *Hs* Na_v_1.4 (55), (IC_50_ = 21.3 ± 1.0 nM, 40.8 ± 1.8 nM, and 6.2 ± 0.4 nM for *Pt* Na_v_1.4, *Dt* Na_v_1.4, and *Pum* Na_v_1.4, respectively) (Fig. S11e-h). Thus, the ability of the poison frogs to resist STX does not arise from their Na_v_s having some unusual resistance to the toxin (Figs. 4, S11d and Table S4). The resistance of poison frogs to the effects of BTX and STX contrasts with the effects of TTX and is not consistent with the high sensitivity of their Na_v_1.4s to all three toxins. These findings suggest that mechanisms, such as toxin sequestration, may prevent BTX and STX from reaching their target Na_v_s.

## Discussion

Poisonous organisms that use toxins as defensive molecules must avoid autotoxication. Such resistance has been proposed to arise from three strategies: 1) acquisition of target protein toxin resistance mutations, 2) toxin sequestration, and 3) enhanced detoxification or elimination capacity (13, 14). Support for the first mechanism includes prominent examples of TTX-resistant Na_v_s in toxin-bearing species and their predators (18–22), STX-resistant Na_v_s in mollusks (17), and epibatidine-resistant nAChRs in poison frogs (16). Because of these examples and the suggestion that the Na_v_ DIVS6 N→T mutation might confer BTX-resistance to poison frogs (25), we expected to find toxin resistant mutants in poison bird and frog Na_v_s. Instead, we found multiple lines of evidence demonstrating that Na_v_s from both poison birds and frogs are highly sensitive to BTX and lack the DIVS6 N→T change. Further, even though the DIVS6 N→T mutation alters the BTX-responses of bird, human, and rat Na_v_1.4s (Figs. 2a-d and S6a-d), it failed to have any effect on the BTX-sensitivity of poison frog Na_v_s (Fig. 2e-h), a result that highlights the importance of vetting putative toxin-resistance mutations in the context of the native channel (16).

How amino acid changes compensate for mutations that alter function is complex and can arise from effects at positions that are far apart in the protein structure (56–59). There are >90 amino acid differences between the poison frog and human Na_v_1.4s (Fig. S4, Table S2) and it is not obvious which variants in the frog Na_v_s suppress the ability of DIVS6 N→T to affect BTX responses. The importance of context is further evident from the fact that even though the Asn position is conserved in all four Na_v_ pore domain subunits, the functional consequences of the N→T change are domain-dependent (Fig. 3, Table S3). These factors, together with the absence of DIVS6 N→T in BTX-bearing birds and frogs (Fig. S1–S2) (24, 26) and its ineffectiveness in poison frog Na_v_s rule out the target-alteration hypothesis for BTX resistance.

Endowing a protein with a new function through mutation often incurs a cost, particularly with respect to protein stability (57–59). Our data show that the DIVS6 N→T change in bird, human, rat, and frog Na_v_s carries substantial functional costs that affect every aspect of channel function, in line with its role in coupling the pore to the voltage-sensor domain in Na_v_s (42), by inducing changes that render the channels more difficult to open, more readily inactivated, and that reduce current density (Table S3), an effect that likely reflects stability penalties that impact channel biogenesis. Similar perturbations of Na_v_ inactivation and reduction of current levels have profound physiological consequences (60) and are linked to a variety of channelopathies (61), underscoring the organism-level fitness problems incurred by changes in Na_v_ biophysical properties. These substantial fitness costs, as well as the inability of the DIVS6 N→T mutation to affect the BTX responses of poison frog Na_v_s are consistent with the low frequency of this variant in wild *P. terribilis* (26) and its absence from the BTX-bearing *P. aurotaenia* poison frog (24). Other studies of ion channel toxin resistance mutants have uncovered various degrees of functional costs that may be compensated by amino acid changes at additional sites in the channel (16, 62). Hence, the effectiveness of developing a toxin-resistant channel via mutation is highly dependent on the cost for evolving this new function and the extent to which functional costs can be mitigated by additional changes.

Poison frogs lacking the Na_v_1.4 DIVS6 N→T change withstand BTX levels that affect non-poisonous frogs (Fig. 4, Table S4), in line with previous studies (23). In non-poisonous frogs, we find clear *in vivo* physiological antagonism between the channel blocker, tricaine, and the channel opener, BTX, that indicates both compounds access the target Na_v_s. This antagonism is absent in poison frogs (Fig. 4, Table S4) even though it can occur at the molecular level of the channel (Fig. S10). Further, the resistance of poison frogs to BTX and STX contrasts with the effects of TTX and is not consistent with the high sensitivity of their Na_v_1.4s to all three toxins (Figs. 4, S10–S11). Together these observations suggest that poison frogs have a means to prevent BTX and STX from reaching their Na_v_s. It is notable that other frogs resist STX poisoning (31, 63, 64) and it is thought that the soluble STX binding protein saxiphilin (13, 31, 65)acts as a ‘toxin sponge’ to sequester and neutralize the lethal effects of this and possibly other neurotoxins (13, 14, 30, 31, 66, 67).

If BTX-bearing animals do not use BTX-resistant Na_v_s to avoid autointoxication, how do they survive? Apart from the absence of BTX-resistant Na_v_s, the diversity among >800 poison frog alkaloid toxins (44), the seasonal and geographical variation of these toxins, and ability to affect multiple ion channels (2) pose major challenges for evolving toxin resistant channels. Enhanced detoxification via metabolic toxin destruction would also not be useful as these poisonous organisms need to handle and store the toxins to deploy them against predators. By contrast, sequestration strategies not only offer a general means of toxin protection, but could act in pathways involved in safely transporting and concentrating toxins in key defensive organs such as the skin (32). The fact that toxin-based chemical defense systems have evolved independently four times in neotropical poison frogs (Dendrobatids)(2), in Malagasy poison frogs (46), and in multiple lineages of poisonous birds (including *Pitohui* and *Ifrita*)(3, 4) supports the idea that such general sequestration mechanisms may underlie toxin autoresistance. Although no BTX-binding proteins have been yet identified, high-affinity toxin binding proteins for STX in frogs, saxiphilin (30, 31, 65) and TTX in pufferfish, Pufferfish Saxitoxin and Tetrodotoxin Binding Protein (68, 69) are known and have been proposed to prevent autointoxication through sequestration (13). Characterizing how such toxin-binding proteins protect hosts from autointoxication, alone or together with specialized toxin transport pathways, should provide new insights into the fundamental mechanisms of toxin autoresistance, expand understanding of how organisms handle a range of chemical insults, and may lead to the discovery of antidotes against various toxic agents.

## Acknowledgements

We thank J. McGlothlin (Virginia Tech) for providing annotated sequences of crow Na_v_s, G. Loussouarn (l’institut du thorax, Nantes) for providing human Na_v_ clones, Z. Chen for technical assistance, and K. Brejc for comments on the manuscript. We thank the Papua New Guinea National Research Institute and Department of Environment and Conservation for permission to conduct fieldwork, the generous people of Bonua Village for assistance in the field permission to work on their land, and the National Geographic Society for funding the fieldwork. This work was supported by grants NIH-NHLBI R01-HL080050 and NIH-NIDCD R01 DC007664 to D.L.M., NSF-1822025 to L.A.O, partial support of this work from NIH-NIGMS GM117263-01A1 to J.D.B, an American Ornithological Society Wetmore Research Award to M.E.K., an NIH-NIGMS F32GM116402 postdoctoral fellowship to R.A.C., and an American Heart Association postdoctoral fellowship to F.A.-A.

## Author Contributions

F.A.-A., M.E.K., R.A.C., C.E.G., J.D.B, J.P.D. and D.L.M. conceived the study and designed the experiments. F.A.-A., N.D.R., R.A.C., and C.E.G. performed patch clamp electrophysiology experiments and analyzed the data. F. A.-A. performed *de novo* Na_v_ cloning, molecular biology experiments, two-electrode voltage clamp electrophysiology experiments, toxin challenge experiments, and analyzed the data. M.E.K. and J.P.D. constructed and analyzed *Pitohui* genomes. J.D.B. synthesized BTX. J.D.B., J.P.D, and D.L.M analyzed data and provided guidance and support. F.A.-A., L.A.O., J.D.B., J.P.D, and D.L.M. wrote the paper.

## Competing interests

J.D.B. is a cofounder and holds equity shares in SiteOne Therapeutics, Inc., a start-up company interested in developing subtype-selective modulators of sodium channels.

The other authors declare no competing interests.

## Data and materials availability

Sequences of *Pum* Na_v_1.4, *Pum* Na_v_1.5, *Pum* Na_v_β2, *Pt* Na_v_1.4, and *Dt* Na_v_1.4 will be deposited with NCBI and released upon publication.

Requests for material to D.L.M.

## Materials and Methods

### Identification and cloning of *Pitohui* Na_v_1.4, Na_v_1.5, and Na_v_β2

Genomic DNA from *Pitohui uropygialis meridionalis* (Family Oriolidae) blood and tissue was extracted using DNeasy kits (Qiagen) to create whole genome sequence libraries for the poisonous *Pitohui* birds. Tissue samples were collected in 1989 from near the village of Bonua, Central Province, Papua New Guinea (10°08’S by 149°10’30” E), stored in ethanol in the field, and frozen since being in the lab. Two genome sequence libraries were created using Illumina Nextera kits. One library had a target insert size of 500-640, and occupied a full run on the MiSeq genetic analyzer using 300-base paired-end reads, and the second library had a target insert size of 640-709 and occupied a full lane of a HiSeq 2500 in rapid run mode using 150-base paired-end reads.

The MiSeq run returned 16,279,946 paired-end reads. The program BBmerge version 4.0 (DOE: Joint Genome Institute; https://jgi.doe.gov/data-and-tools/bbtools/bb-tools-user-guide/bbmerge-guide/) was used to join the forward and reverse reads into a single long read. 7,050,488 of the read pairs (~43%) were joined, and the remaining reads were retained as paired-end reads or single reads for later analyses. The average size of merged reads was 542.6 bases. All reads were then trimmed using Trimmomatic (70) for minimum length, removing adapters, and performing basic quality filtering. All unmerged and unpaired reads were combined into a single fastq file.

The HiSeq run returned 150,979,291 paired-end reads. We removed adapters, trimmed for minimum length, and performed basic quality filtering using Trimmomatic (70). 130,607,604 pairs of reads (86.51%) passed filtering, plus another 9,199,536 (6.09%) of forward only reads passed filter, and 3,342,519 (2.21%) of reverse reads only passed filter. These two read sets (the MiSeq and the HiSeq Illumina datasets) composed the data for gene assembly.

Crows, *Corvus brachyrhynchus* and *Corvus cornix*, (Family Corvidae, from Joel McGlothlin, Virginia Tech) were the closest living relatives of *Pitohuis* that had a fully-annotated genome in the sodium channel gene regions. Crow sequences were used as reference sequences for BLAST searches of Pitohui sequences and for assisting with *Pitohui* Na_v_ assembly. These sequences included the complete nuclear DNA sequence, with all exons, introns, and upstream and downstream UTRs.

To assemble the SCN4A gene from our *Pitohui* reads, we used the following pipeline, written as a bash shell script: First, we used BLATq version 1.0.2 (https://github.com/calacademy-research/BLATq) which uses BLAT (Kent WJ. 2002. BLAT—The BLAST-Like Alignment Tool. Genome Research 12:656-664. DOI: 10.1101/gr.229202. Kent WJ. 2012. BLAT—The BLAST-Like Alignment Tool. Version 35. Available at https://users.soe.ucsc.edu/~kent.) to search both Illumina read sets for any sequences that aligned with the full genome sequences of one of the *Corvus* Na_v_1.4 sodium channel genes. We then used the script excerptByIDs v. 1.0.2 (https://github.com/calacademy-research/excerptByIDs) to create a new fastq files consisting of only the Illumina reads with strong BLATq scores. These files were combined into a single set of all BLAT “hits” using the Unix “cat” command. We then used the assembler SPAdes 3.9.0 (71) to perform an initial *de novo*assembly of reads in the hit file. We improved upon this assembly by using the genome assembly program PRICE ver. 1.2 (72), which iteratively extends the assembly using the assembled contigs from SPAdes as well as all of the existing Illumina read data. See the supplementary materials on the assembly bash shell script and options and parameters.

The assembled contigs from PRICE were loaded into Geneious (ver. 8.0 through 11.0.2). Within Geneious, we used BLAST to identify which contigs contained the *Pitohui* sequences. We created a BLAST database consisting of all of the assembled PRICE contigs, and we used each of the Corvus Na_v_1.4 exons to query the BLAST database of contigs using MegaBLAST. The top hits for each exon suggested which assembled contigs contained the Na_v_1.4 sequence, and typically several to all of the exons were found on the same contig. Assuming that the exon splice patterns were identical in crow and Pitohui, we aligned each of the crow Na_v_1.4 exon sequences to the top-hit Pitohui contig, and annotated the matching exon regions as the Pitohui SCN4A exons.

This assembly and annotation pipelines were repeated for primary voltage-gated sodium channels Na_v_1.5 alpha subunit (SCN5A) as well as the Na_v_βsubunit (SCN2B).

### Cloning of poison frog Na_v_s

Skeletal muscle was harvested from captive *Phyllobates terribilis* and *Dendrobates tinctorius*(Josh’s frogs, Owosso, MI, USA) following euthanasia in accordance with UCSF IACUC protocol AN136799. Total RNA and total DNA were extracted using TRIzol™ Reagent (Thermo Fisher Scientific). Total RNA was reverse transcribed into cDNA using SuperScript™ III First-Strand Synthesis System (Thermo Fisher Scientific). 5’ and 3’ end sequences of genes encoding for *P. terribilis* and *D. tinctorius* Na_v_1.4 were determined by DNA gel extraction and sequencing after rapid amplification of cDNA ends (RACE) using the SMARTer® RACE 5’/3’ Kit (Takara Bio, USA) and internal primers designed from *P. terribilis* and *D. tinctorius* Na_v_1.4 S6 segment sequences (24). From these 5’ end and 3’ end sequences, new primers were designed from both 5’ and 3’ untranslated regions (UTR) of each gene, and were used to amplify full-length *P. terribilis* and *D. tinctorius* Na_v_1.4 genes by polymerase chain reaction (PCR) using Phusion® HF (New England Biolabs). PCR products were gel extracted and sequenced to determine the full-length *P. terribilis*and *D. tinctorius* Na_v_1.4 gene sequences. Direct cloning of the full length PCR products of *P. terribilis* and *D. tinctorius* Na_v_1.4 channel genes into pCDNA3.1 proved problematic resulting unstable constructs prone to deletion. The codon optimized genes were synthesized for expression in human epithelium kidney cells (HEK293) (GenScript, Piscataway, NJ, USA), but were also found to prone to recombination upon insertion into pCDNA3.1. Finally, the gene sequences were redesigned to differ as much as possible from the original genes, synthesized (GenScript, Piscataway, NJ, USA), and cloned into pCDNA3.1. This strategy yielded stable constructs.

### Molecular biology

For electrophysiology experiments, *Hs* Na_v_1.4 (GenBank: NM_000334.4), human Na_v_β1 (GenBank: NM_001037.5), *Pum* Na_v_1.4, *Pum* Na_v_β2, *Pum* Na_v_1.5, *Pt* Na_v_1.4, and *Dt* Na_v_1.4 were subcloned into pCDNA3.1 and *Rn* Na_v_1.4 (GenBank: Y17153.1) was subcloned into pZem228. All mutants were made using the QuikChange Site-Directed Mutagenesis Kit (Agilent) and validated by complete sequencing of the genes encoding for the proteins of interest.

### Patch clamp electrophysiology

Human embryonic kidney cells (HEK 293) were grown at 37 °C under 5% CO2, in a Dulbecco’s modified Eagle’s medium supplemented with 10% fetal bovine serum, 10% L-glutamine, and antibiotics (100 IU ml^−1^ penicillin and 100 mg ml^−1^ streptomycin). HEK293 cells were transfected (in 35-mm-diameter wells) using Lipofectamine 2000 (Invitrogen, Carlsbad, CA, USA) and plated onto coverslips coated with Matrigel (BD Biosciences, San Diego, CA, USA). Human and pitohui Na_v_s were co-expressed with enhanced green fluorescent protein (EGFP), and human Na_v_β1 or *Pitohui* Na_v_β2. Poison frog Na_v_s were co-expressed with EGFP. Transfected cells were identified visually by EGFP expression. A total of 2 μg plasmid DNA (20% Na_v_α, 40% Na_v_β, 40% EGFP) was transfected, except for the poison frog Na_v_s for which a total of 3 μg plasmid DNA (70% Na_v_α, 15% EGFP, 15% SV40 T antigen) was used to increase current amplitude. Experiments studying *Rn* Na_v_1.4 constructs were conducted using Chinese hamster ovary (CHO) cells cultured as described previously (54). Briefly, cells were grown in DMEM (GIBCO, Grand Island, NY) supplemented with 10% cosmic calf serum (HyClone, Logan, UT) and 100 U/mL penicillinstreptomycin (GIBCO). Cells were kept in a 5% CO2 and 96% relative humidity incubator. CHO cells were transfected (in 10 cm plate) using the calcium phosphate precipitation method and EGFP expression as a marker of transfection.

Na^+^ currents were recorded by whole cell patch clamp (73) at room temperature (23 ± 2 °C) 48–72 h post-transfection. Data collection was performed using an Axopatch 200B amplifier (Molecular Devices) and pCLAMP 9 (Molecular Devices, Sunnyvale, CA, USA).

Pipettes were pulled from borosilicate glass capillaries (TW150F-3; World Precision Instruments, Sarasota, FL, USA) and polished with a microforge (MF-900; Narishige, Tokyo, Japan) to obtain 1.2-3.5 MΩ resistances. Sixty to eighty percent of the voltage error due to the series resistance was compensated. For experiments with HEK 293 cells, pipette solution contained the following, in millimolar: 120, Cs methane sulfonate; 8, NaCl; 10, EGTA; 2, Mg-ATP; and 20, Hepes (pH 7.4 with CsOH). Bath solution contained the following, in millimolar: 155, NaCl; 1, CaCl2; 1, MgCl2; 5, KCl; 10, Hepes; and 10, glucose (pH 7.4 with NaOH). For experiments with CHO cells, pipette solution contained the following, in millimolar: 125, CsCl; 40, NaF; 1, EDTA; and 20, Hepes (pH 7.4 with CsOH). Bath solution contained the following, in millimolar: 160, NaCl; 2, CaCl2; and 20, Hepes (pH 7.4 with NaOH).

For experiments with HEK 293 cells, voltage-dependent activation was assessed by stimulating the cells with a multistep depolarization protocol from −90 to +50 mV using 5 mV increments, a −100 mV holding potential, and a sweep-to-weep interval duration of 2 s. Voltage-dependent steady state inactivation was assessed by stimulating the cells with a 500-ms pre-pulse depolarization from −110 to 0 mV in 5 mV steps, followed by a 20-ms step to 0 mV, and repolarization to the holding potential, −100mV; sweep-to-sweep interval duration was 4 s. To examine BTX effects, cells were stimulated upon BTX exposure by applying at least 120 step pulses from −120 to 0 mV at 2 Hz frequency in order to facilitate BTX access into the channel pore as BTX is known to preferentially interact with the open state of Na_v_s (74). For experiments with CHO cells, voltage-dependent activation was assessed by stimulating the cells with a multistep depolarization protocol from −120 to +50 mV using 5 mV increments and a −120 mV holding potential. Voltage-dependent steady state inactivation was assessed by stimulating the cells with a 150-ms pre-pulse depolarization from −140 to 0 mV in 5 mV steps, followed by a 50 ms step to 0 mV, and repolarization to the holding potential, −120mV. Equilibration of BTX was accomplished with persistent activation of channels by applying a 24 ms step depolarization from −120 to 0 mV at a frequency of 2 Hz over the course of 8 min. Leak currents were subtracted using a P/4 protocol during data acquisition. Data analysis was performed using Clampfit 10.6 (Axon Instruments).

Activation curves were obtained by fitting the data with the following single or double Boltzmann equations: I = Imax/(1+exp((V0.5-Vm)/k)), or I = (Imax✓(1+exp((V0.51-Vm)/k1)))+ (Imax2/(1+exp((V0.52-Vm)/k2))), where Imax is the maximal current after normalization to the driving force, V0.5 is the half-activation potential, Vm is the membrane potential, and k is the slope factor. Inactivation curves were obtained by fitting the data with the following single Boltzmann equation: I = Imax/(1+exp((Vm-V0.5)/k)), where Imax is the absolute value of the maximal current at the test pulse, V0.5 is the half-inactivation potential, Vm is the membrane potential, and k is the slope factor. Current density was determined as the ratio between current amplitude (pA) and the membrane capacitance (pF).

### Two-electrode voltage clamp electrophysiology

Two-electrode voltage-clamp recordings were performed on defolliculated stage V–VI *Xenopus laevis* oocytes harvested under UCSF-IACUC protocol AN178461 1-2 days after microinjection with mRNA. Linearized *Pitohui (Pk), P. terribilis (Pt)*, or *D. tinctorius (Dt)* Na_v_1.4 cDNA was translated into capped mRNA using the mMESSAGE mMACHINE™ T7 Transcription Kit (Invitrogen). *Xenopus* oocytes were injected with 0.5-2 ng, 3-6 ng or 10-30 ng of *Pk* Na_v_1.4, *Pt* Na_v_1.4 or *Dt* Na_v_1.4 mRNA, respectively. Two-electrode voltage clamp experiments were performed 1-2 days post-injection. Data were acquired using a GeneClamp 500B amplifier (MDS Analytical Technologies) controlled by pClamp software (Molecular Devices), and digitized at 1 kHz using Digidata 1332A digitizer (MDS Analytical Technologies).

Oocytes were impaled with borosilicate recording microelectrodes (0.3–3.0 MΩresistance) backfilled with 3 M KCl. Sodium currents were recorded using a bath solution containing the following, in millimolar: 96, NaCl; 1, CaCl2; 1, MgCl2; 2, KCl; and 5, HEPES (pH 7.5 with NaOH), supplemented with antibiotics (50 μg ml^−1^ gentamycin, 100 IU ml^−1^ penicillin and 100 μg ml^−1^ streptomycin) and 2.5 mM sodium pyruvate.

For studying the competition between tricaine and BTX, 0.5 mM tricaine was applied by continuous perfusion in the bath solution to assess channel block. BTX was applied from the intracellular side of the membrane by injecting oocytes with 50 nL of 2 mM BTX. Following BTX injection, oocytes were stimulated by applying 1000 step pulses of 60 ms each, from −120 to 0 mV at 2 Hz frequency in order to facilitate BTX access into the channel pore.

To determine STX and TTX dose-response curves, solutions containing test concentrations of each toxin were applied in series by perfusion to oocytes expressing *Pk* Na_v_1.4, *Pt* Na_v_1.4, or *Dt* Na_v_1.4. IC_50_ values were calculated from the ratio of peak currents in the presence and absence of toxin, based on the following equation: 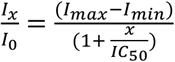 where I_x_ is the current amplitude at the toxin concentration x, I_0_ is the current amplitude in absence of toxin, I_max_ and I_min_ are the maximum and minimum peak current amplitudes, respectively, and IC_50_ is the half-maximal inhibitory concentration.

All toxin effects were assessed with 60-ms depolarization steps from −120 to 0 mV with a holding potential of −120 mV and a sweep-to-sweep duration of 10 s.

Recordings were conducted at room temperature (23 ± 2 °C). Leak currents were subtracted using a P/4 protocol during data acquisition. Data Analysis was performed using Clampfit 10.6 (Axon Instruments) and a custom software developed in the Igor environment (Wavemetrics).

### Toxin challenge experiments

Frogs for the toxin challenge experiments were obtained from the following sources: *Polypedates leucomystax, Phyllobates terribilis*, and *Dendrobates tinctorius* (Josh’s frogs, Owosso, MI, USA); *Xenopus laevis* (Nasco); and *Mantella aurantiaca* (Indoor Ecosystems, Whitehouse, OH, USA). All experiments were performed in accordance with UCSF IACUC protocol AN136799.

Frogs were held at room temperature (23 ± 2 °C) and anesthetized with a 0.15% tricaine (MS-222) bath in prior to toxin injections. Once under anesthesia, as judged by immobility and lack of response to foot pinching, frogs were weighed in order to calculate the appropriate amount of toxin to be administered to at 20x the LD_50_ based on the values calculated for mice as follows: BTX: 2μg per kg, (39), STX: 10 μg per kg (75), and TTX: 12.5 μg per kg, (76). BTX, STX, and TTX, were delivered 40 ng, 200 ng, and 250 ng of toxin, respectively, per g of animal weight. The upper right hind leg was injected with either control, phosphate-buffered saline (PBS), or toxin containing solutions under an SMZ645 binocular microscope (Nikon) using a 30G PrecisionGlideTM needle (BD, Franklin Lakes, NJ, USA). Batrachotoxin (BTX), saxitoxin (STX) or tetrodotoxin (TTX) dissolved in PBS were injected at the appropriate concentration to deliver 20 times the LD_50_. The total volume of injection was 100 μL in *X. laevis* and 10 μL in other frogs due to their smaller size. The choice of intramuscular injection was to avoid internal organ damage. Frogs were allowed to recover in a separate container and monitored constantly for signs of recovery, paralysis, or other adverse symptoms. For *X. laevis*, the recovery container was filled with deionized water and inclined in a way that the frogs could recover on the dry surface of the container base. Post-recovery activity was then assessed by the ability of the frogs to move from the dry to the water-containing surface as *X. laevis* are primarily aquatic animals. For all other frogs, post-recovery activity was assessed by monitoring the ability of the animals to put themselves right-side up from a supine position in their recovery containers. The monitoring period was up to 24 hours post injection.

## Supplementary Materials

**Figure S1.**
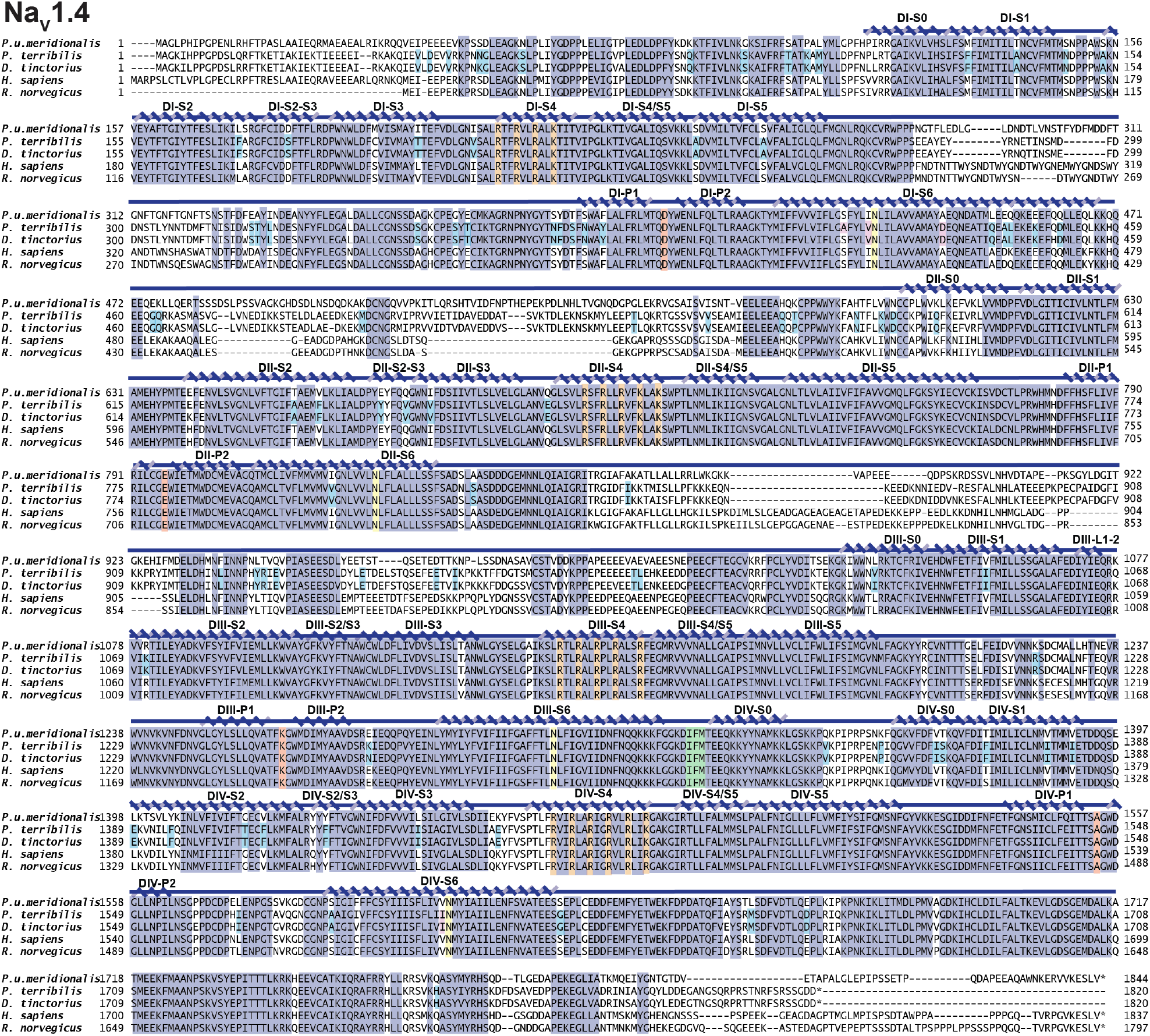
*Pitohui* and Poison frog Na_v_1.4 sequences. Sequence alignment of *Pitohui uropygialis meridionalis* Na_v_1.4, *Phyllobates terribilis* Na_v_1.4, *Dendrobates tinctorius* Na_v_1.4, *Homo sapiens* Na_v_1.4 (NP_000325.4), and *Rattus norgevicus* Na_v_1.4 (NP_037310.1). Key Na_v_ features are highlighted as follows: Selectivity filter ‘DEKA’ (red), ‘IFM’ peptide (green), conserved S6 Asn (yellow), S4 voltage sensor arginines (orange), poison frog variants (cyan) and sites highlighted by (24) (magenta) are indicated. Conserved residues are highlighted (dark blue). Secondary structure elements labeled using boundaries from (77).

**Figure S2.**
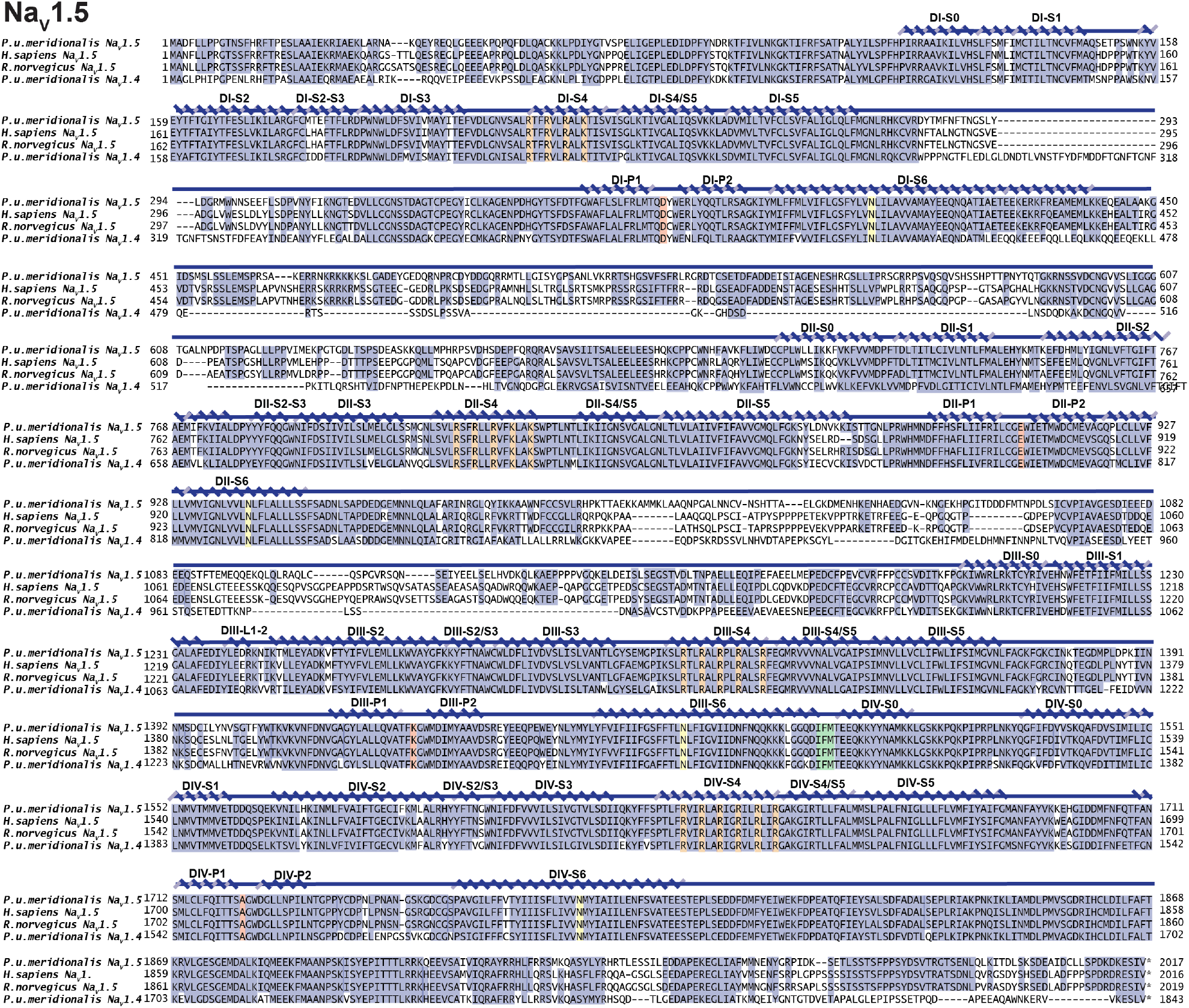
*Pitohui* Na_v_1.5 sequence. Sequence alignment of *Pitohui uropygialis meridionalis* Na_v_1.5, *Homo sapiens* Na_v_1.5 (NP_932173.1), *Rattus norgevicus* Na_v_1.5 (NP_037257.1) *and Pitohui uropygialis meridionalis* Na_v_1.4. Key Na_v_ features are highlighted as follows: Selectivity filter ‘DEKA’ (red), ‘IFM’ peptide (green), conserved S6 Asn (yellow), and S4 voltage sensor arginines (orange). Conserved residues are highlighted (dark blue). Secondary structure elements labeled using boundaries from (77).

**Figure S3.**
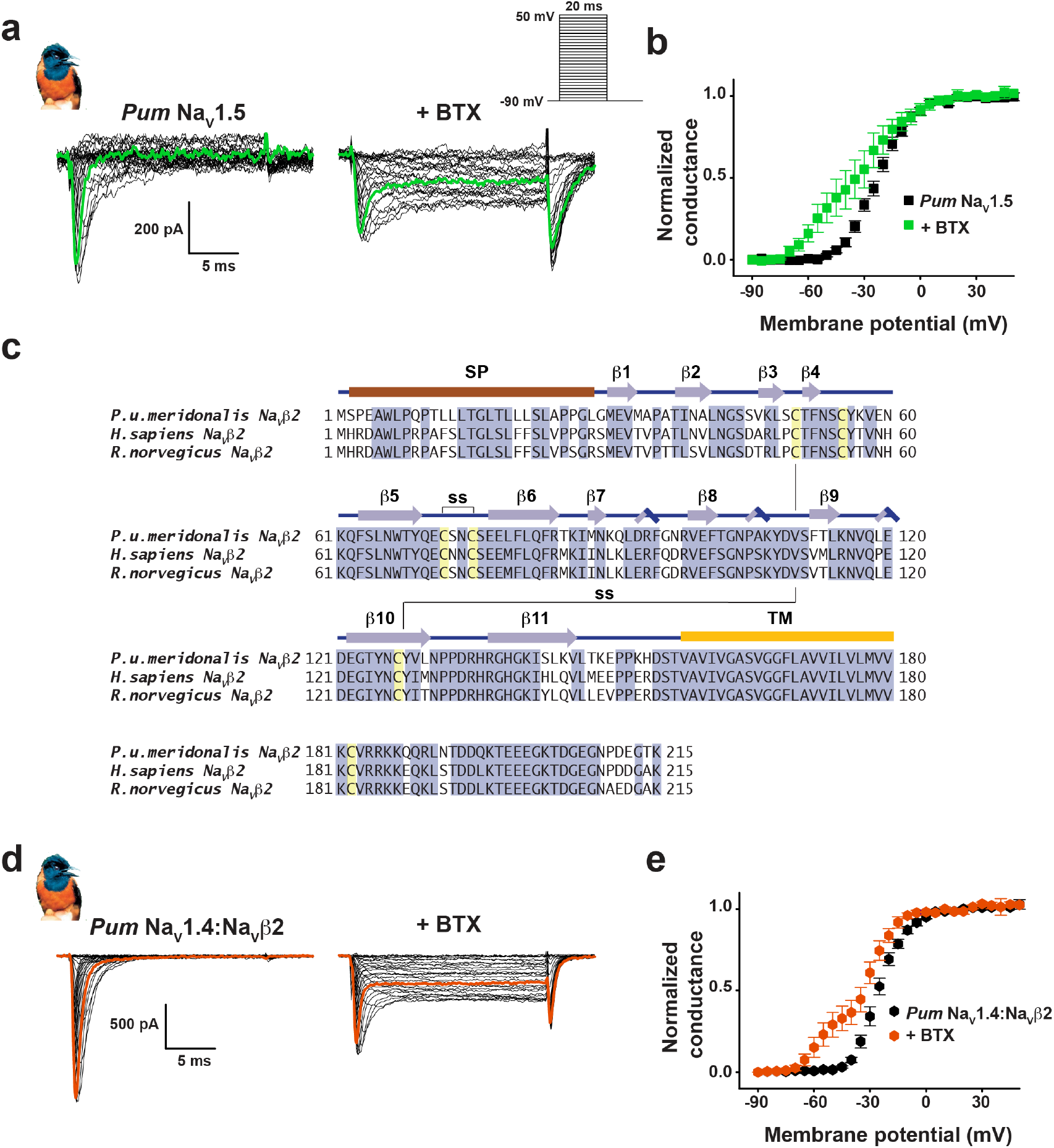
*Pitohui* Na_v_1.5 and Na_v_1.4:Na_v_β2 complexes are BTX sensitive. **a**, Exemplar current recordings for *Pum* Na_v_1.5 expressed in HEK293 cells in absence (left) or presence of 10 μM BTX (right). Trace at 0 mV is highlighted in each panel. Currents were evoked with the shown multistep depolarization protocol (inset). **b**, Conductance-voltage relationships in absence (black squares) or presence of 10 μM BTX (green squares). **c**,Sequence alignment of Na_v_β2 from *Pitohui uropygialis meridionalis, Homo sapiens* (NP_004579.1) and *Rattus norgevicus*(NP_037009.1). Signal peptide (SP), secondary structural elements from (38), conserved disulfide bond (ss), and transmembrane domain (TM) are indicated. **d**, Exemplar current recordings for *Pum* Na_v_1.4:Na_v_β2 expressed in HEK293 cells in absence (left) or presence of 10 μM BTX (right). Trace at 0 mV is highlighted in each panel. Currents were evoked with the shown multistep depolarization protocol (inset panel ‘**a**’). **e**, Conductance-voltage relationships in absence (black hexagons) or presence of 10 μM BTX (red hexagons).

**Figure S4.**
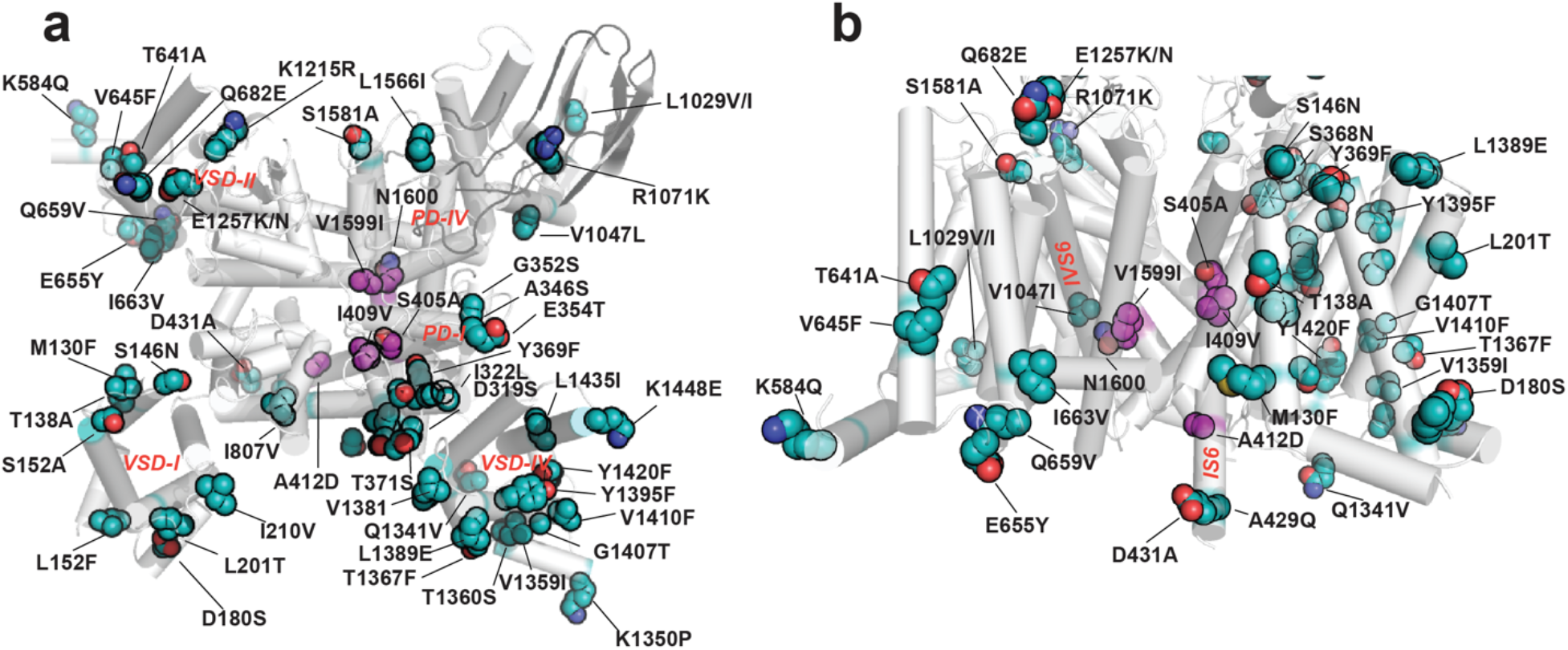
Structural context for poison frog Na_v_ amino acid changes. **a**, and **b**, Locations of poison frog Na_v_ amino acid variants reported here (cyan) and shared with (24) (magenta). Variants are denoted human residue:residue number:frog variant using *Pt* Na_v_1.4 numbering from Fig. S1. Residues are mapped on human Na_v_1.4 (PDB:6ADF) (43). Na_v_1.4 (white).

**Figure S5.**
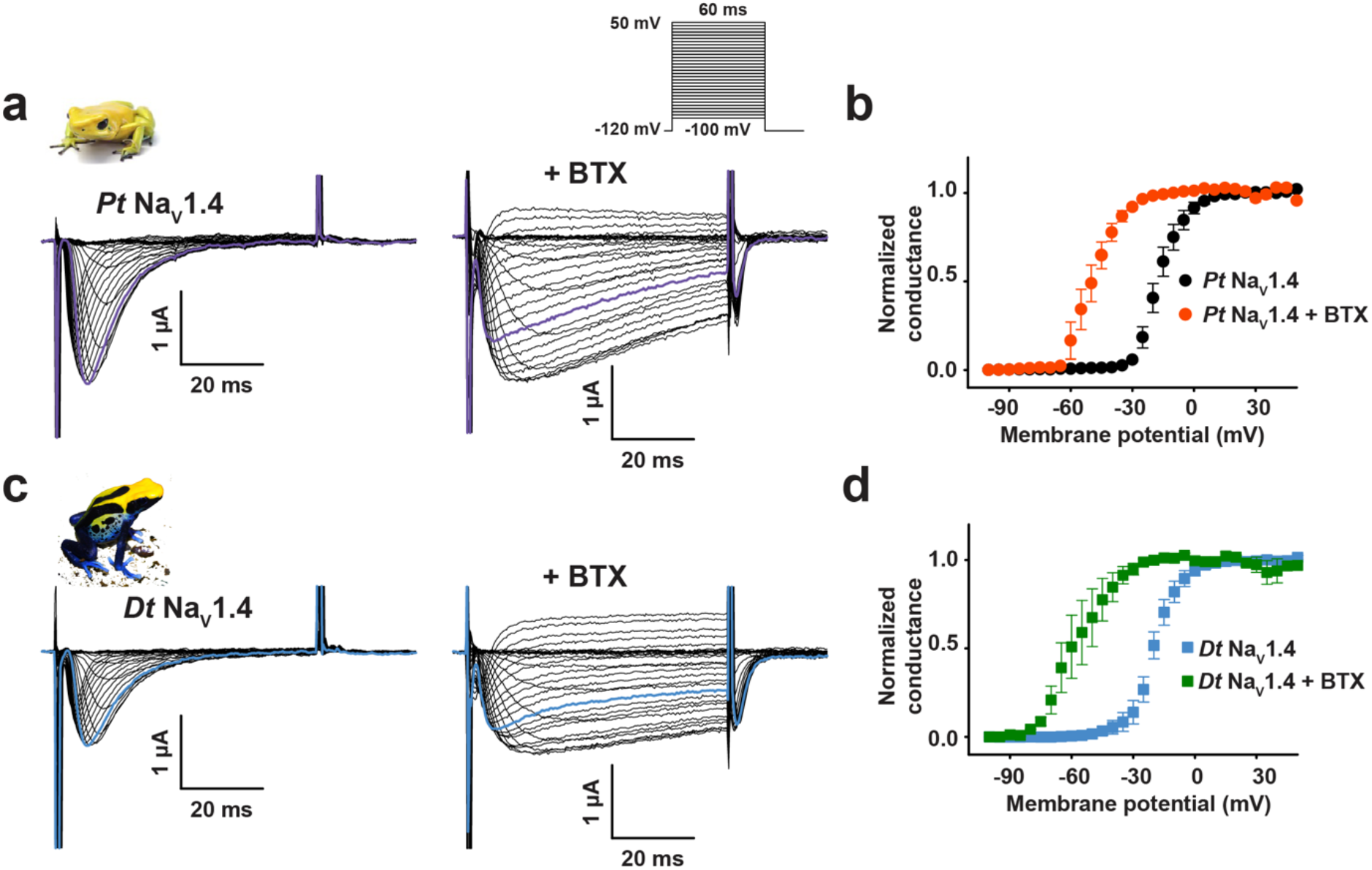
Poison frog Na_v_1.4s expressed in *Xenopus* oocytes are BTX sensitive. Exemplar current recordings (**a** and **c**) for: **a**, *Phyllobates terribilis* Na_v_1.4 (*Pt* Na_v_1.4); and **c**, *Dendrobates tinctorius* Na_v_1.4 (*Dt* Na_v_1.4) expressed in *Xenopus* oocytes in absence (left) or presence of 10 μM BTX (right). Trace at 0 mV is highlighted in each panel. Currents were evoked with the shown multistep depolarization protocol (inset panel ‘**a**’). Conductance-voltage relationships (**b** and **d**) in presence or absence of 10 μM BTX for **b**, *Pt* Na_v_1.4 (black circles), +BTX (orange circles) and **d**, *Dt* Na_v_1.4 (light blue squares), +BTX (green squares).

**Figure S6.**
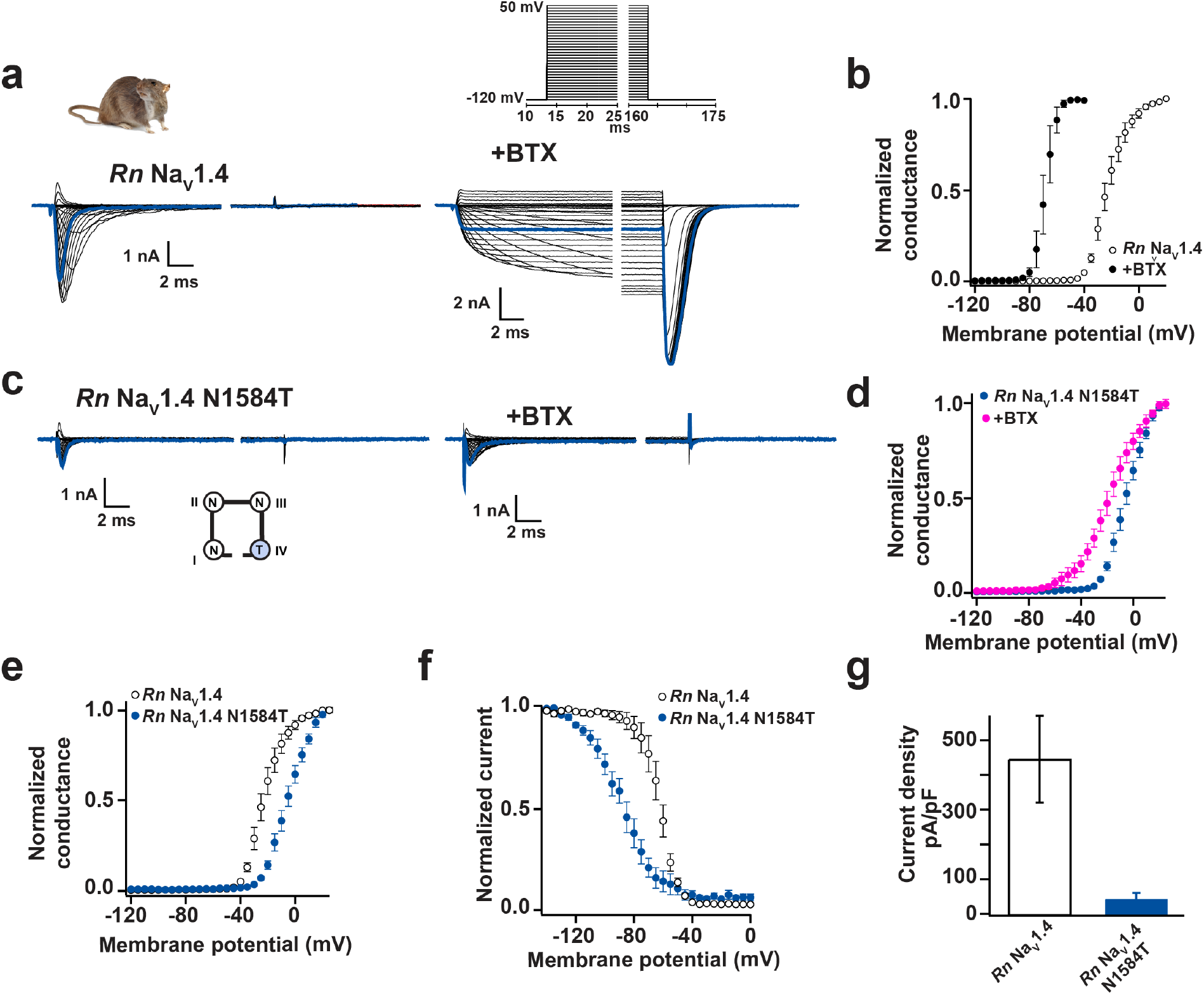
Functional costs of DIV-S6 ‘Asn’ mutation in *Rn* Na_v_1.4. Exemplar current recordings (**a** and **c**) for: **a**, *Rn* Na_v_1.4 and **c**, *Rn* Na_v_1.4 N1584T expressed in CHO cells in absence (left) or presence of 10 μM BTX (right). Trace at 0 mV is highlighted in each panel. Currents were evoked with the shown multistep depolarization protocol (inset panel ‘**a**’). Conductance-voltage relationships (**b** and **d**) in presence or absence of 10 μM BTX for **b**, *Rn* Na_v_1.4 (open circles), +BTX (black circles) and **d**, *Rn* Na_v_1.4 N1584T (blue circles), +BTX (magenta circles). **e**,Conductance-voltage relationships, **f**,Steady-state inactivation voltage dependencies for *Rn* Na_v_1.4 (open circles) and *Rn* Na_v_1.4 N1584T (blue circles). **g**, Current densities for *Rn* Na_v_1.4 (white) and *Rn* Na_v_1.4 N1584T (blue).

**Figure S7.**
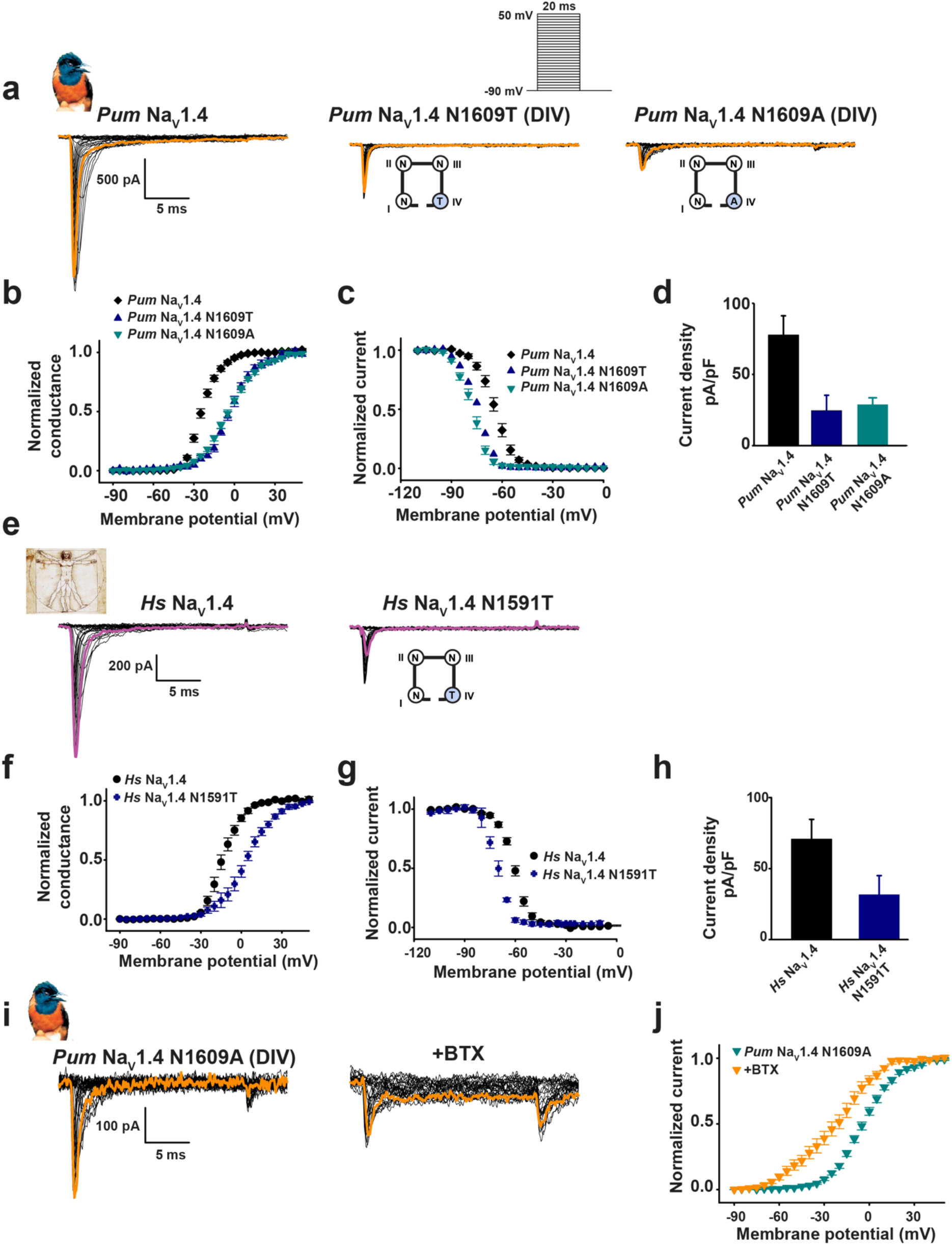
Functional cost of DIV-S6 ‘Asn’ mutation in *Pum* Na_v_1.4 and *Hs* Na_v_1.4. Exemplar current recordings (for: **a**, *Pum* Na_v_1.4 (left), *Pum* Na_v_1.4 N1609T (middle), and *Pum* Na_v_1.4 N1609A (right) expressed in HEK293 cells. Trace at 0 mV is highlighted and currents were evoked with the shown multistep depolarization protocol (inset). Cartoon shows a diagram of the identities of the S6 ‘Asn’ for the ‘Asn’ mutants. **b**,Conductance-voltage relationships, **c**,steady-state inactivation voltage dependencies and **d**, current densities for *Pum* Na_v_1.4 (black diamonds), *Pum* Na_v_1.4 N1609T (blue triangles), and *Pum* Na_v_1.4 N1609A (teal inverted triangles). **e**, Exemplar current recordings for *Hs* Na_v_1.4 (left), *Hs* Na_v_1.4 N1591T (right), expressed in HEK293 cells. Trace at 0 mV is highlighted. Currents were evoked with the shown multistep depolarization protocol from ‘**a**’. **f**,Conductance-voltage relationships, **g**,steady-state inactivation voltage dependencies and **h**, current densities *Hs* Na_v_1.4 (black circles) and *Hs* Na_v_1.4 N1591T (blue diamonds). **i**, Exemplar current recordings for *Pum* Na_v_1.4 N1609A (left) and in the presence of 10 μM BTX (right). **j**, Conductance-voltage relationships for *Pum* Na_v_1.4 N1609A (green inverted triangles) and in the presence of 10 μM BTX (orange inverted triangles).

**Figure S8.**
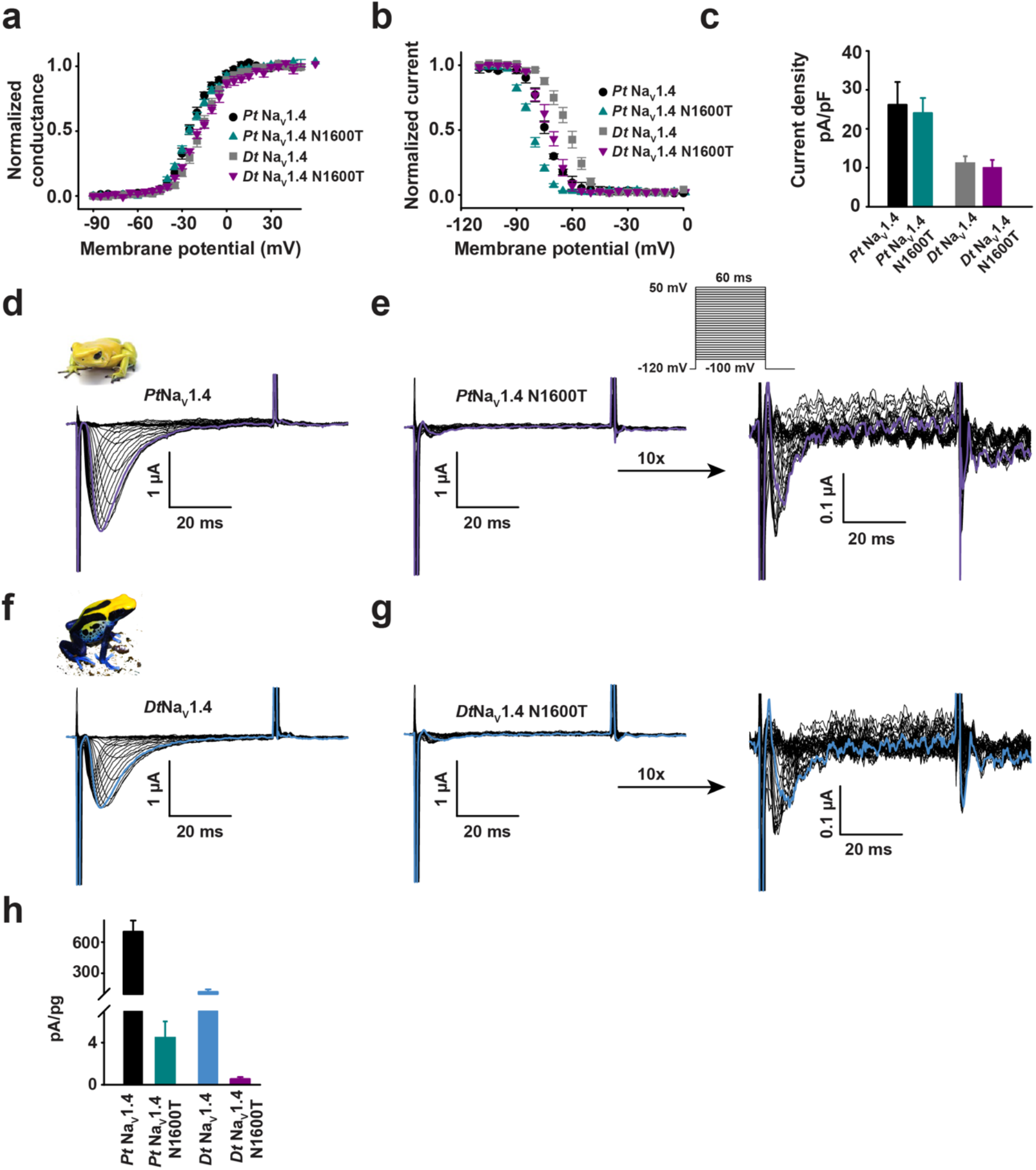
Functional cost of DIV-S6 N→T mutation in poison frog Na_v_1.4s. **a**, Conductance-voltage relationships, **b**, Steady-state inactivation voltage dependences, and **c**, current densities for *Pt* Na_v_1.4, *Pt* Na_v_1.4 N1600T, *Dt* Na_v_1.4, and *Dt* Na_v_1.4 N1600T expressed in HEK293 cells. Exemplar current recordings (**d**, **e**, **f**, and **g**) for: **d**, *Pt* Na_v_1.4, **e**, *Pt* Na_v_1.4 N1600T, **f**, *Dt* Na_v_1.4, and **g**, *Dt* Na_v_1.4 N1600T expressed in *Xenopus* oocytes. 10 times magnification of *Pt* Na_v_1.4 N1600T and *Dt* Na_v_1.4 N1600T traces are shown (**e** and **g**, right panels). Trace at 0 mV is highlighted in each panel. Currents were evoked with the shown multistep depolarization protocol (inset panel ‘**e**’). **h**, Current amplitudes normalized to the amount of injected RNA for the indicated poison frog constructs.

**Figure S9.**
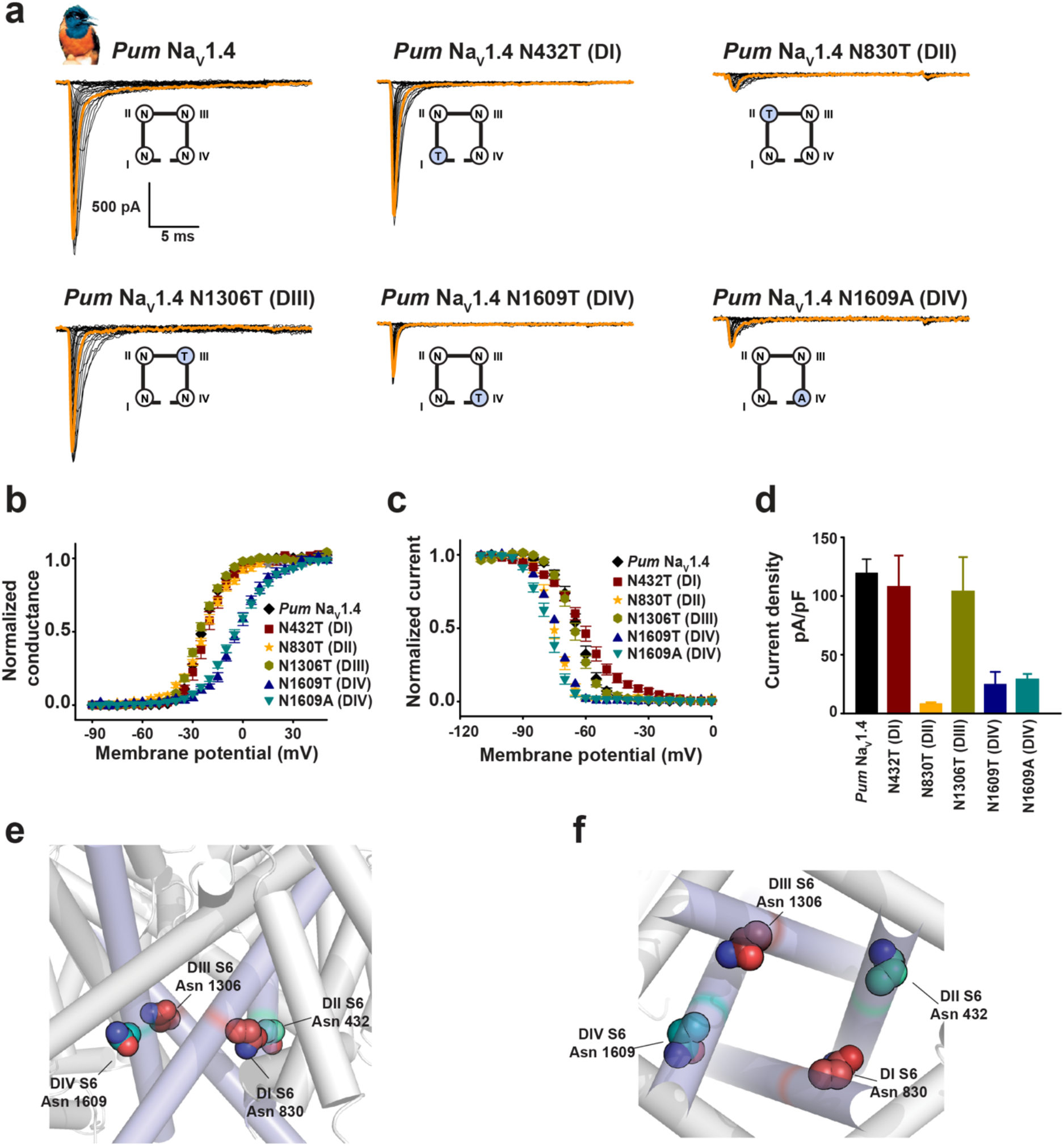
Functional studies of S6 Asn mutants support asymmetric properties of the channel pore. **a**, Exemplar current recordings, **b**, conductance-voltage relationships, **c**, steady-state inactivation voltage dependences, and **d**, current densities for *Pum* Na_v_1.4, *Pum* Na_v_1.4 N432T, *Pum* Na_v_1.4 N830T, *Pum* Na_v_1.4 N1306T, *Pum* Na_v_1.4 N1609T, and *Pum* Na_v_1.4 N1609A expressed in HEK293 cells. Trace at 0 mV is highlighted in each panel. Cartoon shows a diagram of the identities of the S6 ‘Asn’ for each construct. **e**, and **f**,Side ‘e’ and bottom ‘f’ views of the locations the S6 conserved asparagines. Residues are mapped on the structure of human Na_v_1.4 (PDB:6ADF) (43) and are labeled using the *Pum* Na_v_1.4 numbering.

**Figure S10.**
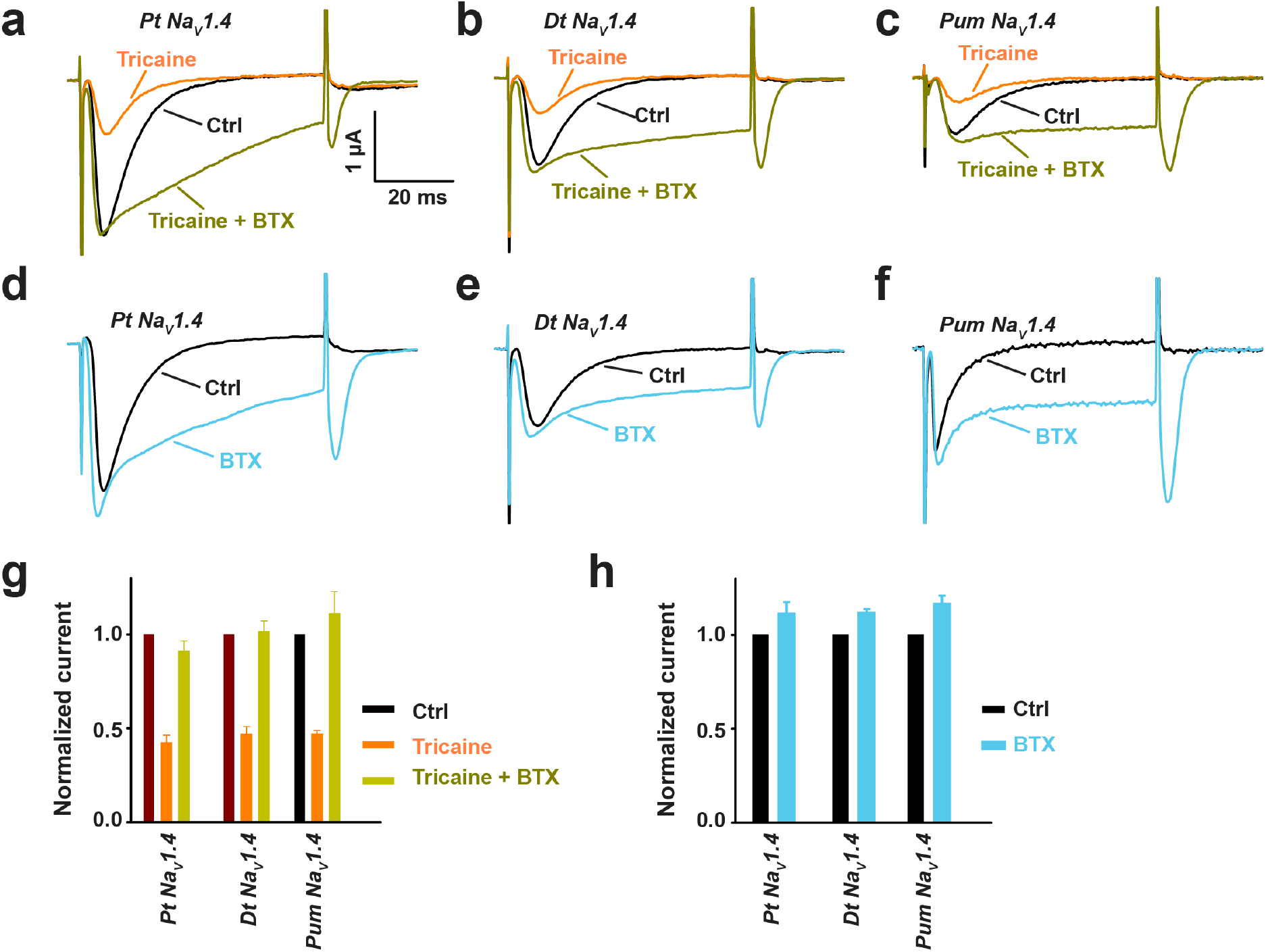
BTX competes with anesthetic agent tricaine in Na_v_s from poisonous species. **a**, Exemplar recordings at 0 mV in control (black), after 0.5 mM tricaine application (orange) and after BTX injection (dark green) into the same *Xenopus* oocytes expressing *Pt* Na_v_1.4 (left), *Dt* Na_v_1.4 (middle) or *Pum* Na_v_1.4 (right). BTX injection was performed after tricaine block of sodium current and the recordings of the BTX effect were done while the oocyte was still exposed to tricaine. **b**, Exemplar recordings at 0 mV before (black) or after BTX injection (light blue) into the same *Xenopus* oocytes expressing Na_v_1.4 from the indicated poisonous species. **c** and **d**, Average peak current amplitudes normalized to the corresponding control peak current amplitude.

**Figure S11.**
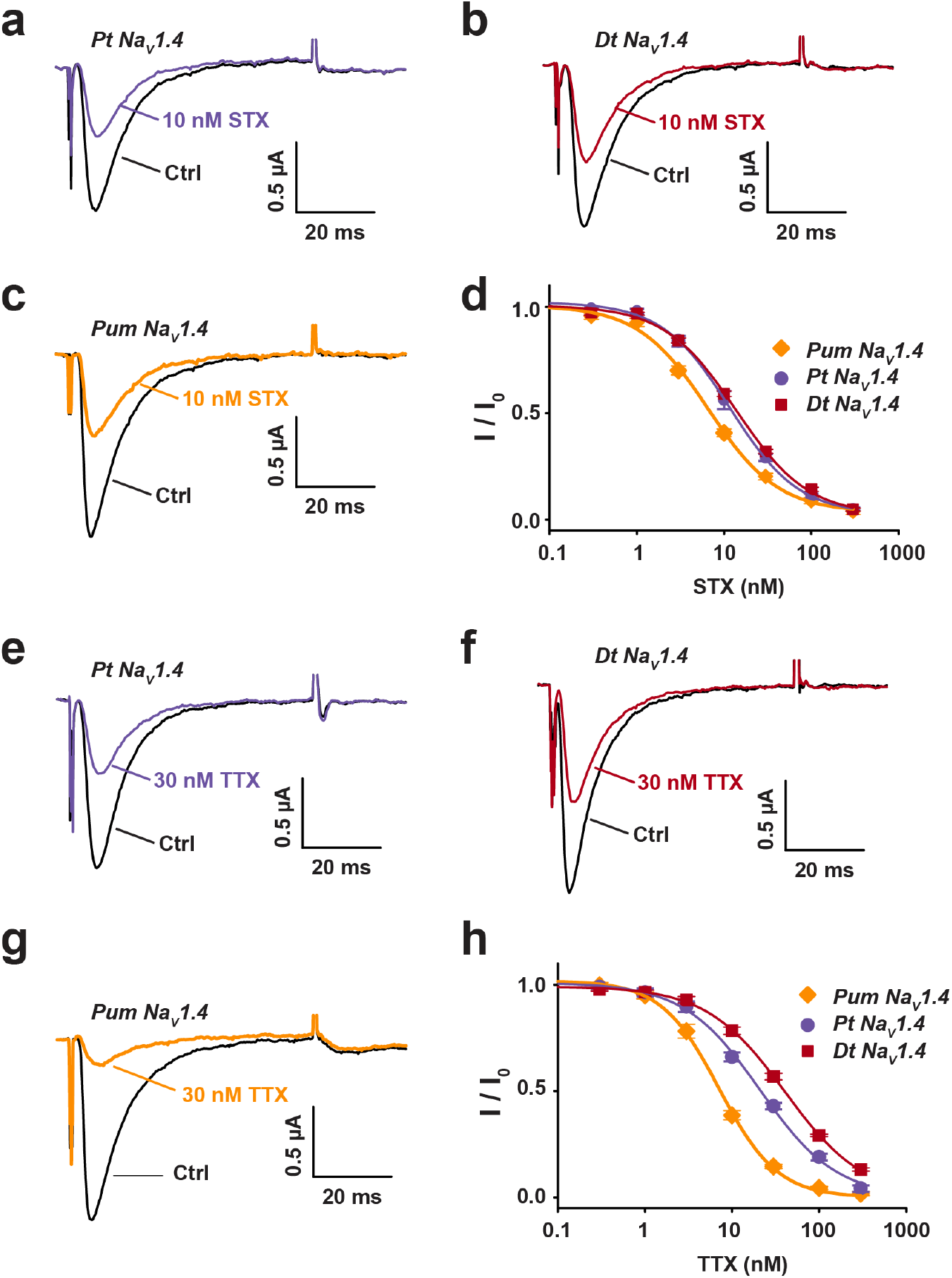
Na_v_1.4s from poisonous animals are STX and TTX sensitive. **a-c**, Exemplar recordings at 0 mV before (black) and after 10 nM STX application to *Xenopus* oocytes expressing **a**, *Pt* Na_v_1.4 (purple) **b**, *Dt* Na_v_1.4 (red), or **c**, *Pum* Na_v_1.4 (orange). **d**, STX Dose-response curves for *Pt* Na_v_1.4 (purple circles), *Dt* Na_v_1.4 (red squares), and *Pum* Na_v_1.4 (orange diamonds). Curves show fits to the Hill equation. (IC_50_ = 12.6 ± 1.4 nM, 14.6 ± 0.6 nM, and 7.3 ± 0.5 nM for *Pt* Na_v_1.4, *Dt* Na_v_1.4, and *Pum* Na_v_1.4, respectively. Errors are SEM. (n = 4). **e-g**, Exemplar recordings at 0 mV before (black) and after 30 nM TTX application to *Xenopus*oocytes expressing **e**, *Pt* Na_v_1.4 (purple), **f**, *Dt* Na_v_1.4 (red), or **g**, *Pum* Na_v_1.4 (orange). **h**, TTX Dose-response curves for *Pt* Na_v_1.4 (purple circles), *Dt* Na_v_1.4 (red squares), and *Pum* Na_v_1.4 (orange diamonds). Curves show fits to the Hill equation. (IC_50_ = 21.3 ± 1.0 nM, 40.8 ± 1.8 nM, and 6.2 ± 0.4 nM for *Pt* Na_v_1.4, *Dt* Na_v_1.4, and *Pum* Na_v_1.4, respectively. Errors are SEM. (n = 5-6).

**Table S1.** Na_v_ Inactivation parameters

| Channel |  | V<sub>1/2 inact</sub> | ΔV<sub>1/2 inact</sub> | k<sub>inact</sub> | n |
| --- | --- | --- | --- | --- | --- |
| **Bird** | ***Pum* Na<sub>v</sub>1.4** | -64.2 ±1.3 | n.a. | 4.8 ±0.1 | 14 |
|  | ***Pum* Na<sub>v</sub>1.4 + *Pk*Na<sub>v</sub>β2** | -66.5 ±1.4 | n.a. | 4.6 ±0.2 | 8 |
|  | ***Pum* Na<sub>v</sub>1.5** | -75.9 ±1.6 | n.a. | 4.2 ±0.5 | 6 |
|  | ***Pum* Na<sub>v</sub>1.4 N432T (DI)** | -61.2 ±1.6 | n.a. | 9.8 ±0.5 | 6 |
|  | ***Pum* Na<sub>v</sub>1.4 N830T (DII)** | -75.0 ±0.9 | -10.8 ±1.6 | 4.2 ±0.2 | 7 |
|  | ***Pum* Na<sub>v</sub>1.4 N1306T (DIII)** | -65.3 ±1.0 | n.a. | 5.1 ±0.5 | 7 |
|  | ***Pum* Na<sub>v</sub>1.4 N1609T (DIV)** | -74.5 ±1.7 | -10.3 ±2.1 | 4.9 ±0.1 | 3 |
|  | ***Pum* Na<sub>v</sub>1.4 N1609A (DIV)** | -77.6 ±0.9 | -13.4 ±1.6 | 4.7 ±0.2 | 10 |
| **Human** | ***Hs* Na<sub>v</sub>1.4** | -60.5 ±0.8 | n.a. | 4.7 ±0.3 | 12 |
|  | ***Hs* Na<sub>v</sub>1.4 N1591T (DIV)** | -70.3 ±0.9 | -9.8 ±1.2 | 4.2 ±0.2 | 3 |
| **Rat** | ***Rn* Na<sub>v</sub>1.4** | -61.9 ±0.2 | n.a. | 6.1 ±0.2 | 4 |
|  | ***Rn* Na<sub>v</sub>1.4 N1584T (DIV)** | -88.9 ±0.4 | -27.0 ±0.4 | 12.6 ±0.4 | 6 |
| **Poison frog** | ***Pt* Na<sub>v</sub>1.4** | -73.9 ±0.6 | n.a. | 5.9 ±1.4 | 3 |
|  | ***Pt* Na<sub>v</sub>1.4 (N1600T) (DIV)** | -81.6 ±0.7 | -7.7 ±0.9 | 4.7 ±0.3 | 6 |
|  | ***Dt* Na<sub>v</sub>1.4** | -62.2 ±1.3 | n.a. | 5.6 ±0.5 | 8 |
|  | ***Dt* Na<sub>v</sub>1.4 (N1600T) (DIV)** | -72.0 ±1.4 | -9.8 ±1.9 | 5.2 ±0.3 | 3 |

**Table S2.**
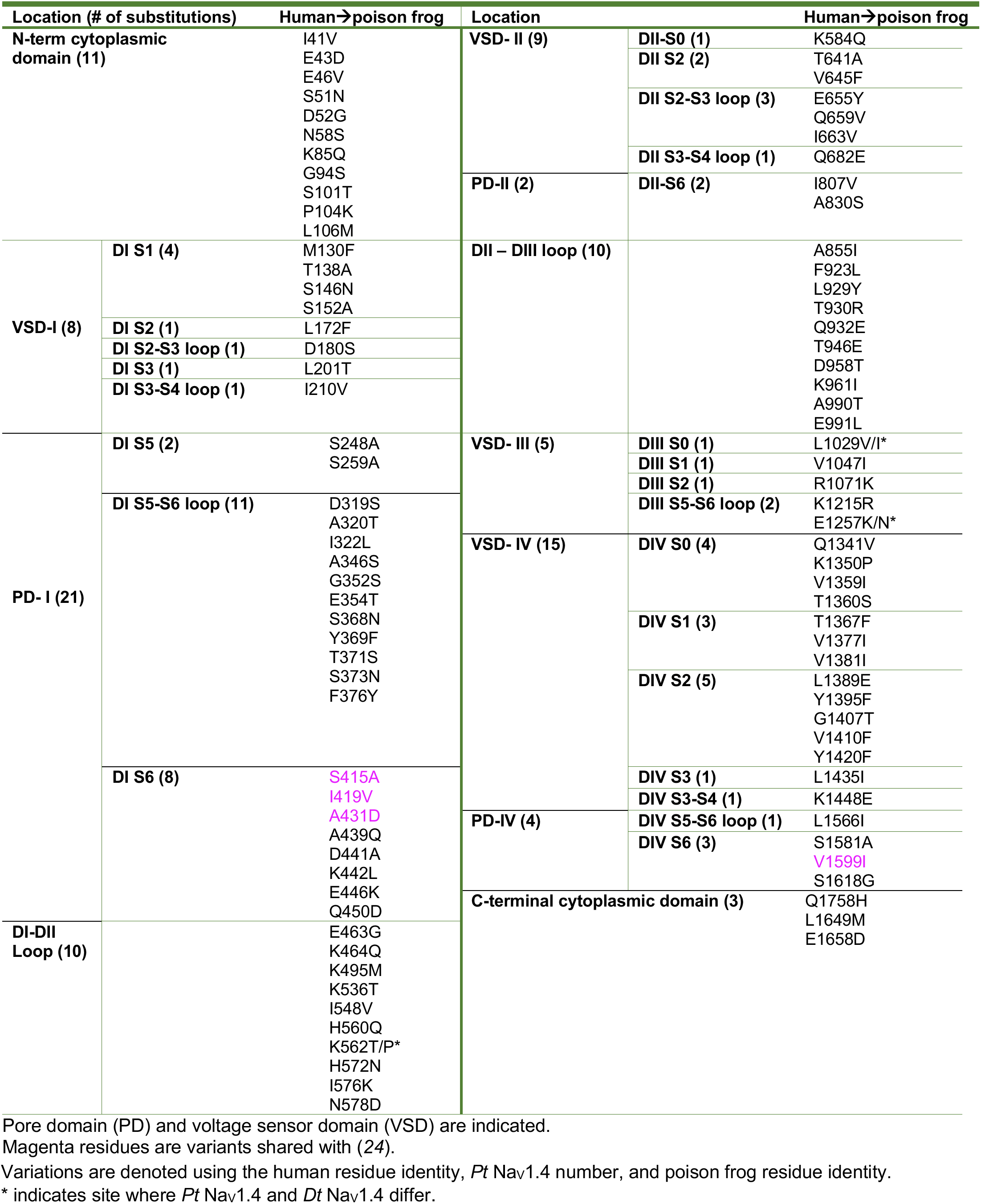
Human→Poison frog Na_v_1.4 amino acid variants

**Table S3.** Comparison of the functional effects of S6 mutations

| Channel | BTX resistant? | Activation shift?<br>$\Delta V_{1/2 \text{ act}}$ (mV) | Inactivation shift?<br>$\Delta V_{1/2 \text{ inact}}$ (mV) | Decrease in current density?<br>(fold change) |
| --- | --- | --- | --- | --- |
| *Pum* Nav1.4 N432T (DI) | No | No | No | No |
| *Pum* Nav1.4 N830T (DII) | No | No | Yes; -10.8 ± 1.6 | Yes; 14.8 |
| *Pum* Nav1.4 N1306T (DIII) | No | No | No | No |
| *Pum* Nav1.4 N1609T (DIV) | Yes | Yes; +20.5 ± 1.5 | Yes; -10.3 ± 2.1 | Yes; 4.9 |
| *Pum* Nav1.4 N1609A (DIV) | No | Yes; +19.6 ± 1.6 | Yes; -13.4 ± 1.6 | Yes; 4.1 |
| *Hs* Nav1.4 N1591T (DIV) | Yes | Yes; +16.9 ± 2.7 | Yes; -9.8 ± 1.2 | Yes; 2.2 |
| *Rn* Nav1.4 N1584T (DIV) | Yes | Yes; +18.0 ± 0.6 | Yes; -27.0 ± 0.4 | Yes; 10.3 |
| *Pt* Nav1.4 (N1600T) (DIV) | No | No | Yes; -7.7 ± 0.9 | No* |
| *Dt* Nav1.4 (N1600T) (DIV) | No | No | Yes; -9.8 ± 1.9 | No* |

Data represent measurements in transfected HEK293 cells
'\*' indicates substantial current density loss in *Xenopus* oocytes.

**Table S4.** Recovery time from anesthesia (minutes)

|  | **PBS** | **BTX** | **STX** | **TTX** |
| --- | --- | --- | --- | --- |
| ***X. laevis*** | 29 ± 1 | 15 ± 2 | N/A | N/A |
| ***P.leucomystax*** | 169 ± 12 | 70 ± 20 | N/A | 720 ± 30 |
| ***P. terribilis*** | 10 ± 1 | 14 ± 2 | 72 ± 8 | 354 ± 6 |
| ***D. tinctorius*** | 35 ± 4 | 38 ± 3 | 32 ± 14 | 283 ± 57 |
| ***M. aurantiaca*** | 56 ± 11 | 60 ± 10 | 52 ± 3 | 1011 ± 51 |

Values are average ± S.E.M.

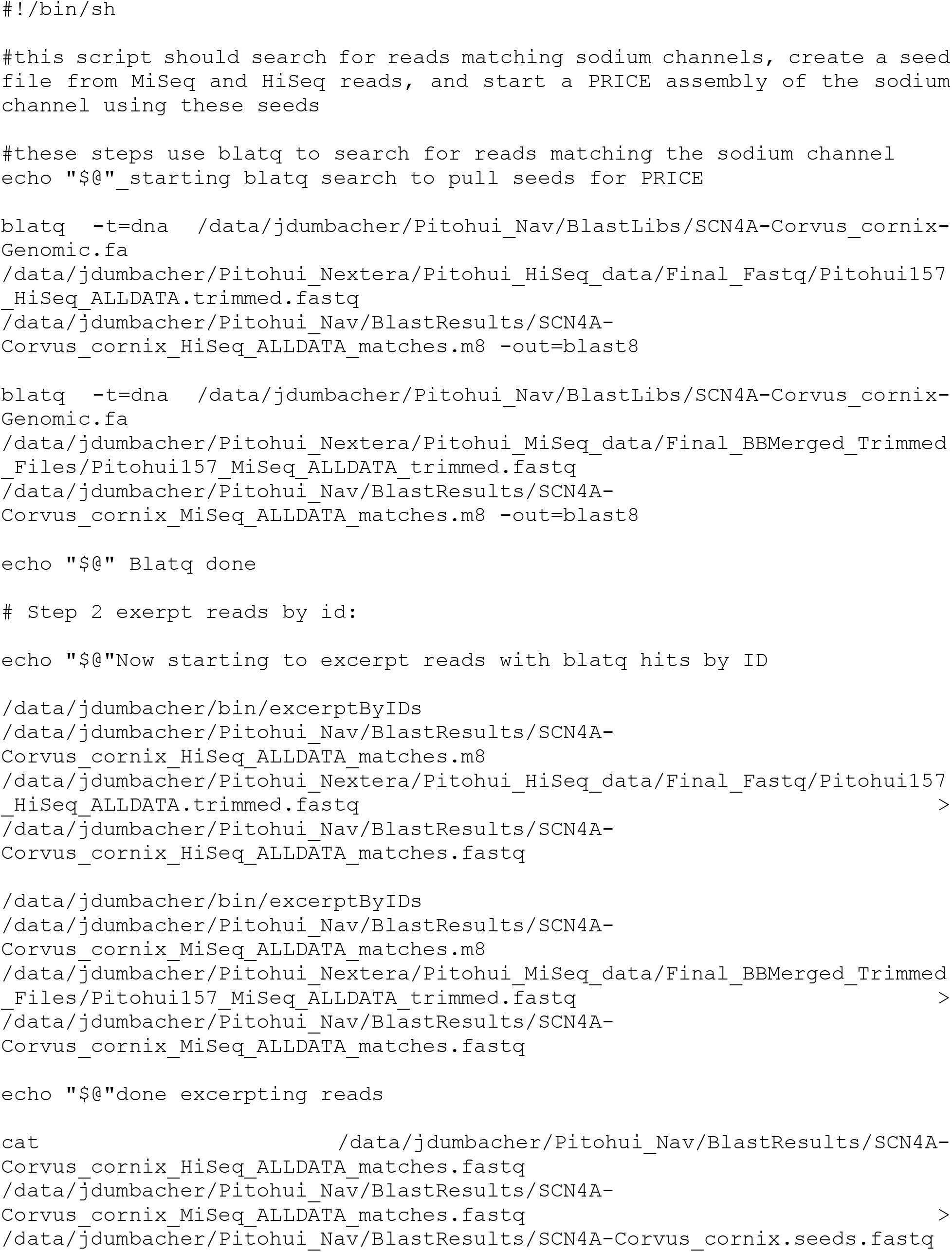
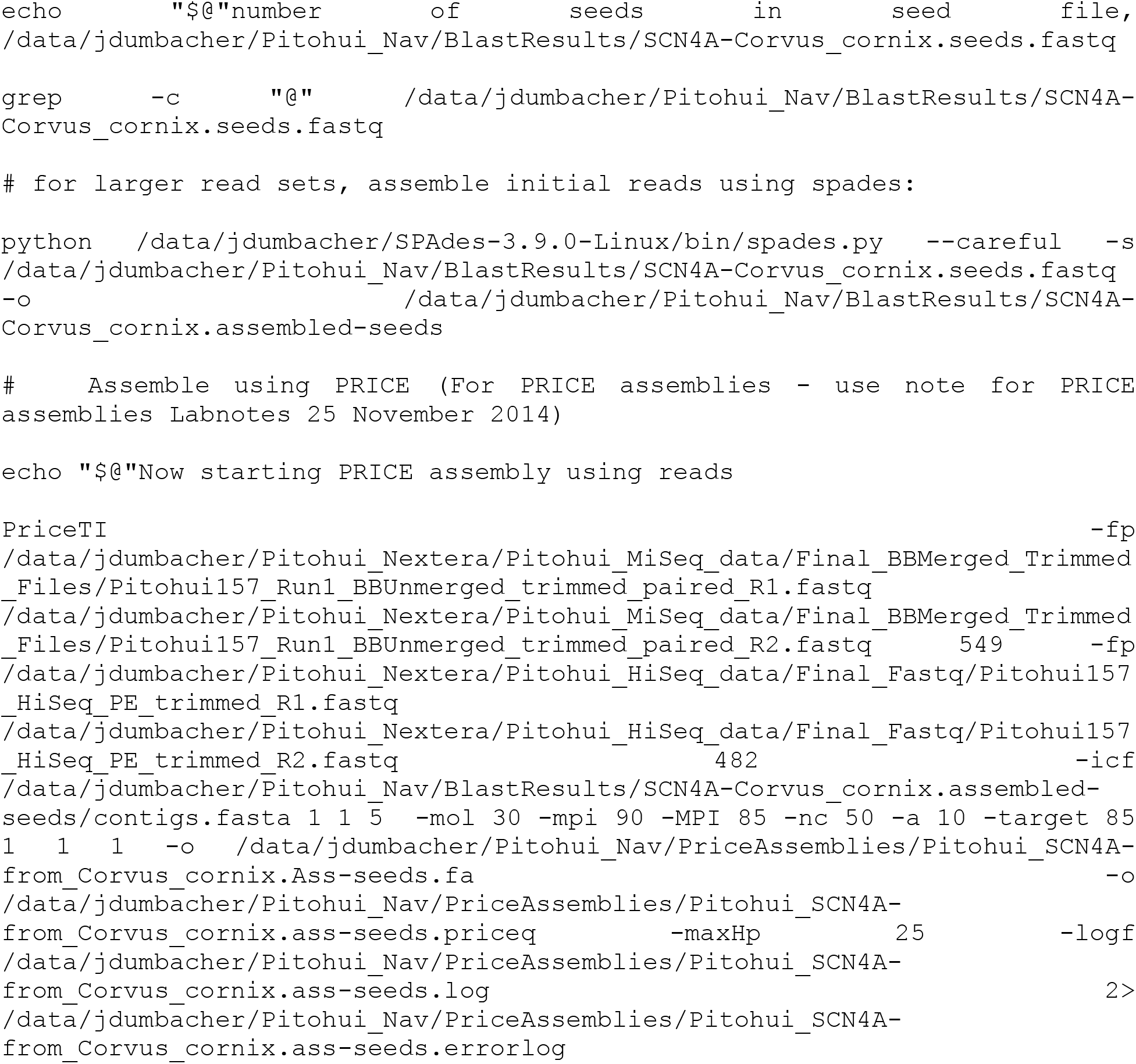
Gene assembly scripts.

